# Ketone body driven lipid remodeling supports thermogenic adaptation to fasting

**DOI:** 10.1101/2025.06.13.659540

**Authors:** Zahra Moazzami, Eric D. Queathem, Jiawen Ma, Qian Chen, Rini Arianti, Ifrah Aden, Anneliese Brown, Zamzam Abdulahi, Chenxin Gu, Xuechun Chang, Yin Li, Haidong Yu, Alessandro Bartolomucci, Satchidananda Panda, Douglas G. Mashek, Endre Kristóf, Patrycja Puchalska, Lisa S. Chow, Zhihao Jia, Peter A. Crawford, Hai-Bin Ruan

## Abstract

Metabolic adaptation to fasting may have conferred survival advantage to early humans and predicts weight gain caused by overnutrition in modern societies. Fasting suppresses brown adipose tissue (BAT) thermogenesis; however, it is unclear how BAT rewires cellular metabolism to balance between energy conservation and heat generation. Here, we report that BAT in mice under fasting and cold challenge consumed ketone bodies, specifically acetoacetate (AcAc). Ablating liver ketogenesis decreased, while enhancing hepatic AcAc output defended, body temperature in mice facing the dual challenge. Using stable isotope tracing in brown adipocytes in vitro combined with quantitative analysis of metabolic fluxes and lipidomics in BAT from genetic mouse models, we disentangled the two metabolic fates of AcAc – terminal oxidation in the mitochondria and lipid biosynthesis in the cytosol. Notably, AcAc-sourced carbon preferentially supported polyunsaturated fatty acid synthesis in BAT, linking to the positive impact of intermittent fasting on lipid profiles in both mice and humans. Therefore, ketone body utilization by thermogenic adipocytes contributes to metabolic resilience of mammals and can be targeted to optimize benefits of dietary regimens.

## Introduction

Food availability is a strong evolutionary pressure that has shaped the metabolic traits and physiological adaptations of all animals including humans. During the transition between feeding and fasting states, inter-organ communication and coordination enable the body’s flexibility to metabolize different energy sources, such as carbohydrates and fat (Bartolomucci and Speakman; Castillo-Armengol et al., 2019; Priest and Tontonoz, 2019; Shulman and Petersen, 2012). Conversely, metabolic inflexibility is considered as a hallmark and mechanism of obesity and related disorders (Goodpaster and Sparks, 2017). Fasting induces a profound switch in fuel utilization from glucose to fatty acids and fatty acid-derived ketone bodies, which is evolutionarily conserved in mammals for them to survive periods of food scarcity. In modern societies facing a global obesity epidemic, fasting regimens such as intermittent fasting (IF) and time-restricted feeding (TRF) have been suggested as potential preventive strategies to improve metabolic inflexibility and reduce inflammation (Marko et al., 2024). Importantly, fasting-induced energy expenditure responses not only determine weight loss during fasting but also predict weight gain during overfeeding (Hollstein et al., 2019). However, our understanding of thermogenic adaptation to fasting remains limited.

Brown adipose tissue (BAT) is a specialized thermogenic organ that mammals have evolved for cold adaptation (Cannon and Nedergaard, 2004). BAT dissipates chemical energy as heat through mitochondrial uncoupling protein 1 (UCP1) (Keipert et al., 2024), as well as UCP1-independent mechanisms (Roesler and Kazak, 2020). Beyond thermogenesis, BAT also serves as a “metabolic sink” for excessive nutrients and secretes “batokines” to regulate systemic metabolism (Bartelt et al., 2011; Harms and Seale, 2013; Schulz and Tseng, 2013; Yoneshiro et al., 2019). In humans, the prevalence of BAT is inversely associated with body mass index (Betz and Enerback, 2011; Tam et al., 2012), age (Saito et al., 2009; van Marken Lichtenbelt and Schrauwen, 2011; Yoneshiro et al., 2011), and incidence of cardiometabolic diseases (Becher et al., 2021). Along with the great interest in the relationship between BAT and energy balance, there are debates in the field regarding whether enhancing BAT might be a therapeutic target to treat cardiometabolic diseases (Cypess et al., 2025; Gupta, 2023). A possible avenue to resolve the dispute is to study BAT metabolism in conditions resembling selective pressure that shapes thermogenic adaptation.

During winter’s cold temperatures, animals often also face food scarcity. To survive such extreme environments, should non-hibernating animals activate BAT thermogenesis to stay warm or restrain BAT activity for energy conservation (Reinisch et al., 2020; Ruan et al., 2014)? In hibernating mammals that have been fasted for months, what energy fuels BAT thermogenesis to initiate arousals from hypothermic torpor (Ballinger and Andrews, 2018)? Answers to these questions remain open, partially because prior research on BAT fuel utilization was predominantly performed in animals that were fed *ad libitum*. In these conditions, cold stimulates the utilization of fatty acids, acylcarnitines, glucose, lactate, and amino acids in BAT; however, it remains undermined if these nutrients are equally importantly during fasting.

The thermogenic and endocrine function of BAT requires the dynamic remodeling of the lipid composition of adipocytes, which is perturbed by obesity and aging (Cho et al., 2023). Recent lipidomics-based approaches revealed that thermogenic activation induces the glycerophospholipid pathway in BAT, leading to increased abundance of phosphatidylglycerol (PG) and phatidylethanolamine (PE), particularly those containing long-chain and unsaturated fatty acids in their acyl chains (Lynes et al., 2018; Marcher et al., 2015; Rajakumari and Srivastava, 2021). Phospholipid remodeling supports mitochondrial biogenesis and thermogenic metabolism of BAT, and also importantly, fuels the synthesis of bioactive “lipokines” to modulate systemic physiology (Tsuji and Tseng, 2023). Nonetheless, the fuel sources and molecular regulation of BAT lipid remodeling have been less characterized.

Here, we discovered that BAT metabolizes liver-produced ketone bodies for fasting-induced thermogenic adaptation and energy conservation in mice. BAT preferentially utilizes acetoacetate (AcAc) over D-β-hydroxybutyrate (D-βOHB) for both uncoupling thermogenesis and shunting into fatty acid biosynthesis and phospholipid remodeling. Ketone metabolism in thermogenic adipocytes provides survival advantages in extreme environments and mediates the metabolic benefits of fasting, suggesting potential therapeutic applications for treating lipid abnormalities in cardiometabolic diseases.

## Results

### Hepatic ketogenesis supports thermogenic adaptation to fasting

Compared to mice fed *ad libitum* (AL) at room temperature (RT), 16h of overnight fasting followed by 6h of cold exposure at 4 °C reduced core body temperature (**Fig. S1A**) and interscapular BAT (iBAT) temperature (**Fig. 1A**). To investigate changes in BAT fuel sourcing imposed by fasting and cold, we collected arterial blood from the left ventricle and venous blood from the Sulzer’s vein draining iBAT and determined the arterio-venous (AV) differences in metabolite concentrations (**Fig. 1B**) (Park et al., 2023). Log2 ratios of A-to-V blood concentration were calculated to quantify relative uptake (positive values) or release (negative values) of each metabolite. BAT during fasting consumed similar amount of glucose but less triglycerides (TGs) and amino acids (**Fig. 1C, S1B**). Metabolomics of AV samples revealed no significant changes in lactate or acylcarnitines (**Fig. S1C**). Since fasting induces ketosis (**Fig. S1D**), we examined whether ketone bodies were consumed by BAT in the setting of fasting and cold exposure. Indeed, total ketone bodies (TKB) showed net zero exchange in AL-RT mice but net consumption in the fast-cold condition (**Fig. 1D**). D-β-hydroxybutyrate (D-βOHB) is the major ketone body in the circulation; however, its flux rate in BAT was not affected by fasting (**Fig. 1E**). Instead, BAT showed increased absorption (and/or reduced release) of the other ketone body acetoacetate (AcAc), when mice were challenged with fasting and cold (**Fig. 1F**). These results demonstrate that thermogenic adaptation to fasting is associated with increased AcAc consumption in BAT.

**Figure 1.**
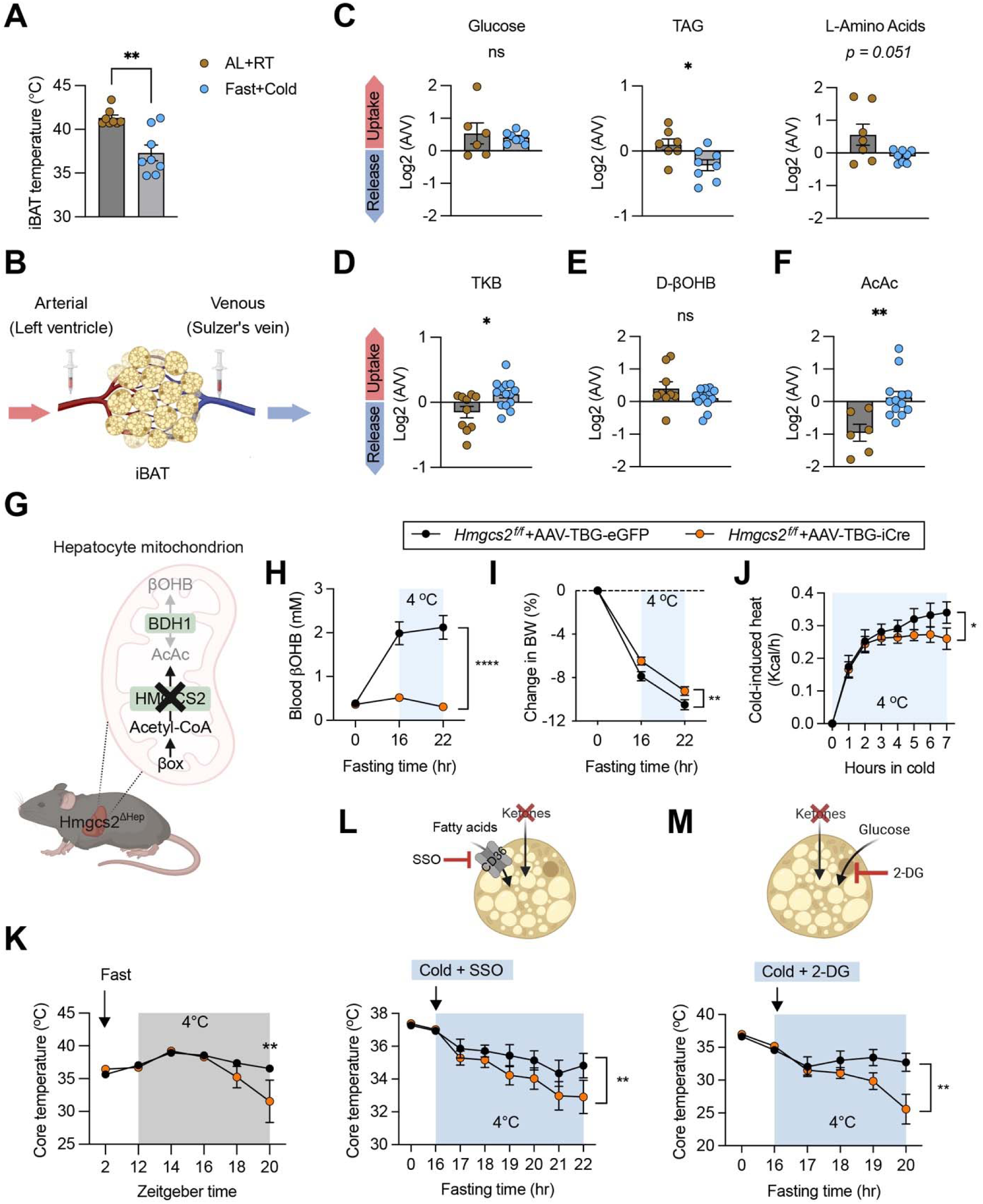
Hepatic ketogenesis for thermogenic adaptation to fasting. (**A**) Interscapular BAT temperature of AL-RT and Fast-Cold mice (n = 8), determined by infrared thermography. (**B-F**) Left ventricle (A) and Sulzer’s vein (V) blood samples from AL-RT and Fast-Cold mice were collected (B) and subjected to measurements of (C) glucose, TG, amino acids, (D) total ketone body (TKB), (E) D-βOHB, and (F) AcAc. (**G-H**) *Hmgcs2*^Δ*Hep*^ mice were generated (G) and subjected to overnight fasting followed with 8 hours of cold challenge. Blood D-βOHB (H), weight loss (I), and energy expenditure (J) were determined. (**K**) *Hmgcs2*^Δ*Hep*^ mice were fasted at ZT2 (light cycle) and exposed to cold at ZT12 (night cycle). Rectal temperature was measured. (**L, M**) Rectal temperature of overnight fasted *Hmgcs2*^Δ*Hep*^ mice that received CD36 inhibitor SSO (L) or glycolysis inhibitor 2-DG (M) at the beginning of cold exposure. Data are represented as mean ± SEM. * P < 0.05, ** P < 0.01, *** P < 0.001 by two-tailed unpaired Student’s t-test (A, C-F) and two-way ANOVA (H-M) and n.s., not significant. Please also see Figure S1.

Ketone bodies are primarily synthesized in the liver, mediated by mitochondrial 3-hydroxymethylglutaryl-CoA synthase (HMGCS2). To determine the role of hepatic ketogenesis in BAT thermogenesis, we injected AAV8 expressing Cre recombinase driven by the TBG promoter (AAV-TBG-iCre) into *Hmgcs2^f/f^*mice to generate *Hmgcs2*^Δ*Hep*^ knockout animals (**Fig. 1G and Fig. S1E**). Compared to AAV-TBG-eGFP injected controls, *Hmgcs2*^Δ*Hep*^ mice were devoid of fasting-induced ketosis (**Fig. 1H**). Reduced weight loss in *Hmgcs2*^Δ*Hep*^ mice suggested a reduction in energy expenditure when ketones were absent (**Fig. 1I**). Indeed, indirect calorimetry revealed a lower increase of cold-induced energy expenditure in fasted *Hmgcs2*^Δ*Hep*^ mice compared to controls (**Fig. 1J**). Attenuated body temperature control could be observed when fasted *Hmgcs2*^Δ*Hep*^ mice were exposed to cold during the dark cycle (**Fig. 1K**), but not evidently during the light cycle (**Fig. S1F**), underscoring the circadian responses to energetic challenges in animals (van der Vinne et al., 2018). Metabolomics quantification of AV samples from these animals revealed reduced release of fatty acids and increased uptake of amino acids by BAT (**Fig. S1G**), indicative of compensation from other nutrients when ketones are absent. To test this, we administered sulfo-N-succinimidyl oleate (SSO), an inhibitor of CD36 that blocks fatty acid uptake, and found that fasted *Hmgcs2*^Δ*Hep*^ mice were more cold-sensitive than *Hmgcs2^f/f^* mice, even during the light cycle (**Fig. 1L**). When glucose utilization was blocked by 2-deoxyglucose (2-DG), more profound decrease in body temperature was observed in *Hmgcs2*^Δ*Hep*^ mice (**Fig. 1M**), suggesting the increased dependence on glucose metabolism when ketones are not available to BAT. These results show that the ketone bodies can serve as alternative fuels for BAT when animals are cold challenged in the setting of prior fasting.

### Acetoacetate (AcAc) flux enhances thermogenic adaptation to fasting

To test if brown adipocytes can directly use ketones, we isolated stromal vascular fraction (SVF) cells from mouse iBAT and differentiated them into primary brown adipocytes. D-βOHB and AcAc were supplemented in physiological concentrations and media were collected to measure consumption (**Fig. 2A, B**). AcAc is less stable than D-βOHB (Puchalska et al., 2021), but when adjusted for stability, we observed a significant uptake of AcAc, but not D-βOHB, by brown adipocytes (**Fig. 2B, C**). This is consistent with the net consumption of AcAc observed in BAT in vivo (**Fig. 1F**). D-βOHB dehydrogenase (BDH1) mediates the interconversion between AcAc and D-βOHB in the mitochondria (Puchalska and Crawford, 2017; Stagg et al., 2021). The inefficient metabolism of D-βOHB might be due to the relatively low expression of BDH1 in BAT. Nonetheless, BDH1 protein could be detected in human BAT and positively correlated with UCP1 expression (**Fig. S2A**) (Muller et al., 2016). In cold exposed mice, fasting modestly increased BDH1 expression in BAT (**Fig. S2B**), suggesting potentially increased ketone interconversion and D-βOHB utilization in such condition.

**Figure 2.**
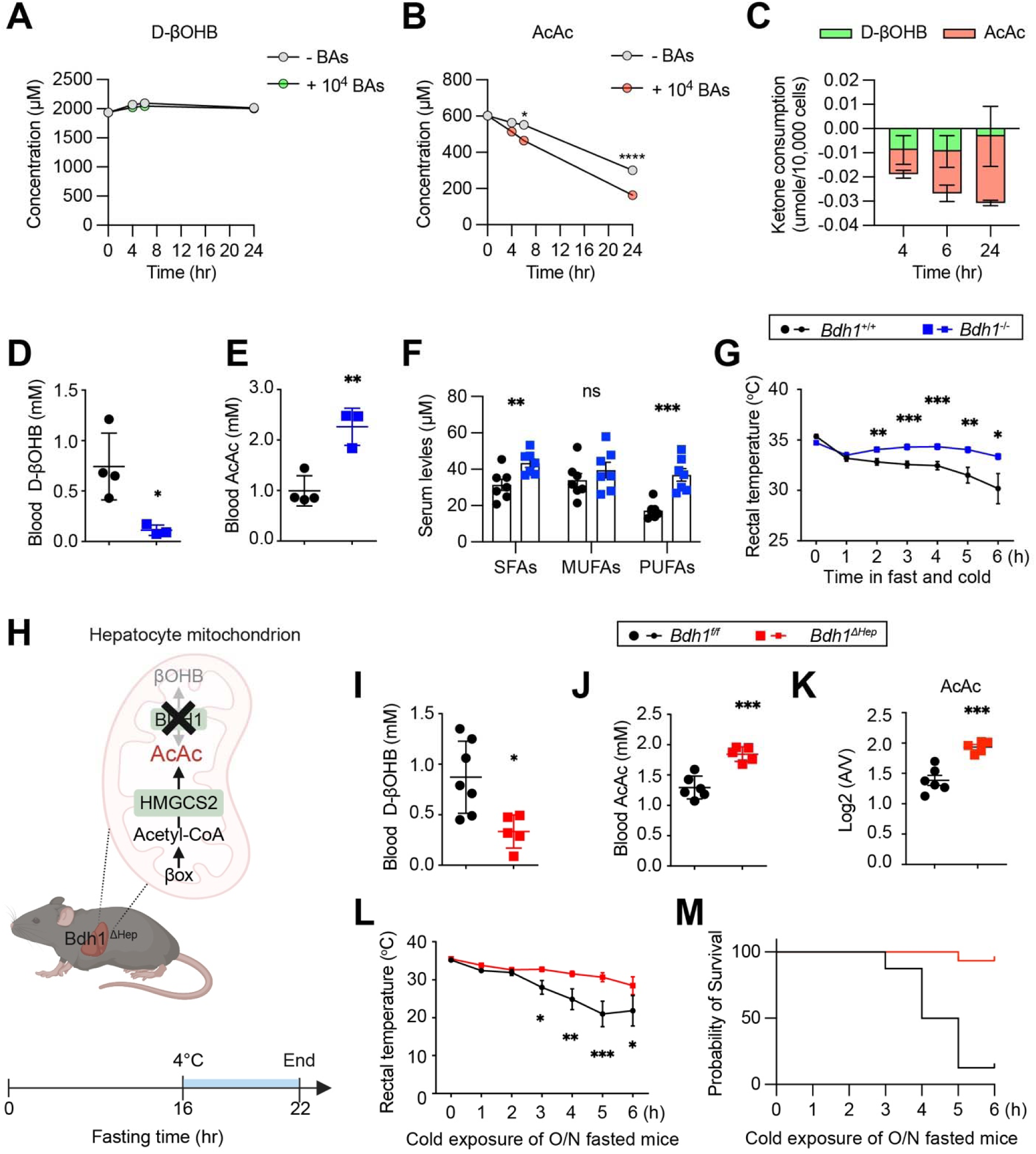
Ketone body aceteoacetate (AcAc) promotes BAT thermogenesis. (**A-C**) Differentiated brown adipocytes (BAs) were supplemented with D-βOHB (A) and AcAc (B) and media were collected at indicated time points for measurement and calculation of ketone body consumption (C). (**D-F**) Serum of overnight fasted *Bdh1*^+/+^ and *Bdh1*^-/-^ mice were measured for D-βOHB (D) and AcAc (E) with a kit and FFAs (F) with lipidomics. (**G**) Core body temperature of *Bdh1*^+/+^ and *Bdh1*^-/-^ mice subjected to simultaneous challenge with fasting and cold. (**H-K**) *Bdh1*^Δ*Hep*^ mice were generated (H) and subjected to overnight fasting. Blood D-βOHB (I), blood AcAc (J), and AcAc consumption by BAT (K) were determined. (**L, M**) Overnight fasted *Bdh1*^Δ*Hep*^ mice and their control were challenged with cold. Rectal temperature (L) and survival curve (M) were shown. Data are represented as mean ± SEM. * P < 0.05, ** P < 0.01, *** P < 0.001 by two-way ANOVA (A, B, G, L), two-tailed unpaired Student’s t-test (D-F, I-K) and Kaplan-Meier Analysis (M). Please also see Figure S2.

Considering that AcAc was the primary ketone used by BAT, we sought to test the physiological outcome of augmented AcAc flux. Global *Bdh1*^-/-^ mice represent a unique model that produces and uses only one ketone body (AcAc) over the other (D-βOHB). As expected, these animals profoundly reduced levels of circulating D-βOHB (**Fig. 2D**) and increased the availability of AcAc systemically (**Fig. 2E**). Supporting the role of AcAc in fueling *de novo* lipogenesis and polyunsaturated fatty acid (PUFA) elongation (Queathem et al., 2025), we observed a doubling of circulating PUFA levels in *Bdh1*^-/-^ mice (**Fig. 2F**). When subjected to concurrent challenge with fasting and cold, *Bdh1*^-/-^ mice were able to maintain a higher body temperature than controls (**Fig. 2G**). RNA-sequencing of BAT revealed that genes upregulated by BDH1 deficiency were involved in endoplasmic reticulum homeostasis, cell-cell junction, and mitochondrial matrix (**Fig. S2C**).

Similarly, liver-specific *Bdh1*^Δ*Hep*^ mice (*Alb-Cre*; *Bdh1^f/f^*) also had higher circulating AcAc (**Fig. 2H-J**), as we previously reported (Stagg *et al*., 2021). Indeed, when measuring AV samples from *Bdh1*^Δ*Hep*^ mice, we observed substantial increase in AcAc absorption (**Fig. 2K**). Consequently, *Bdh1*^Δ*Hep*^ mice were more resilient to fasting and cold-induced hypothermia (**Fig. 2L**) and mortality (**Fig. 2M**). On the other hand, ablating BDH1 only in the skeletal muscle did not interfere systemic equilibration between AcAc and D-βOHB (**Fig. S2D, E**), nor cold tolerance of fasted mice (**Fig. S2F**). Together, these results demonstrate that metabolic flux of AcAc from the liver to BAT promotes adaptive thermogenesis during fasting.

### BAT ketone metabolism is a survival strategy

When supplemented during brown adipogenesis from SVF cells, neither AcAc nor D-βOHB evidently changed adipogenesis (**Fig. S3A**), thermogenic gene expression (**Fig. S3B**), or UCP1 protein expression (**Fig. S3C**). To test if ketones can directly stimulate thermogenic activity, we measured the oxygen consumption rate (OCR) of primary brown adipocytes in glucose-containing medium supplemented with ketone bodies. Mixed AcAc and D-βOHB in physiological 1:3 ratio dose-dependently increased basal, uncoupling, and adrenergic receptor (AR) agonist-stimulated OCR (**Fig. 3A, B and Fig. S3D**). Compared to D-βOHB, AcAc induced much stronger ATP-dependent respiration and proton leak (**Fig. 3C, D**). Increased OCR was not due to redox-dependent interconversion by BDH1, because inverting ratios between AcAc and D-βOHB had similar stimulative effect on OCR (**Fig. S3E**). However, when compared to glucose and palmitate, ketones demonstrated much lower capacity to stimulate OCR of brown adipocytes in a nutrient-deprived medium (**Fig. 3E and Fig. S3F**). Using Pimozide, a selective antagonist against 3-oxoacid CoA-transferase 1 (OXCT1) that mediates ketone catabolism (Al Batran et al., 2020), we found that AR activation by norepinephrine (NE) increased the capacity of brown adipocytes to oxidize ketones (**Fig. 3F**), correspondingly leading to decreased dependence on ketone oxidation (**Fig. 3G**). In comparison, brown adipocytes had the highest capacity of and dependence on glucose oxidation (**Fig. 3F, G**). These data suggest that ketones might be less thermogenic fuels and may intricate with glucose and lipid metabolism in BAT.

**Figure 3.**
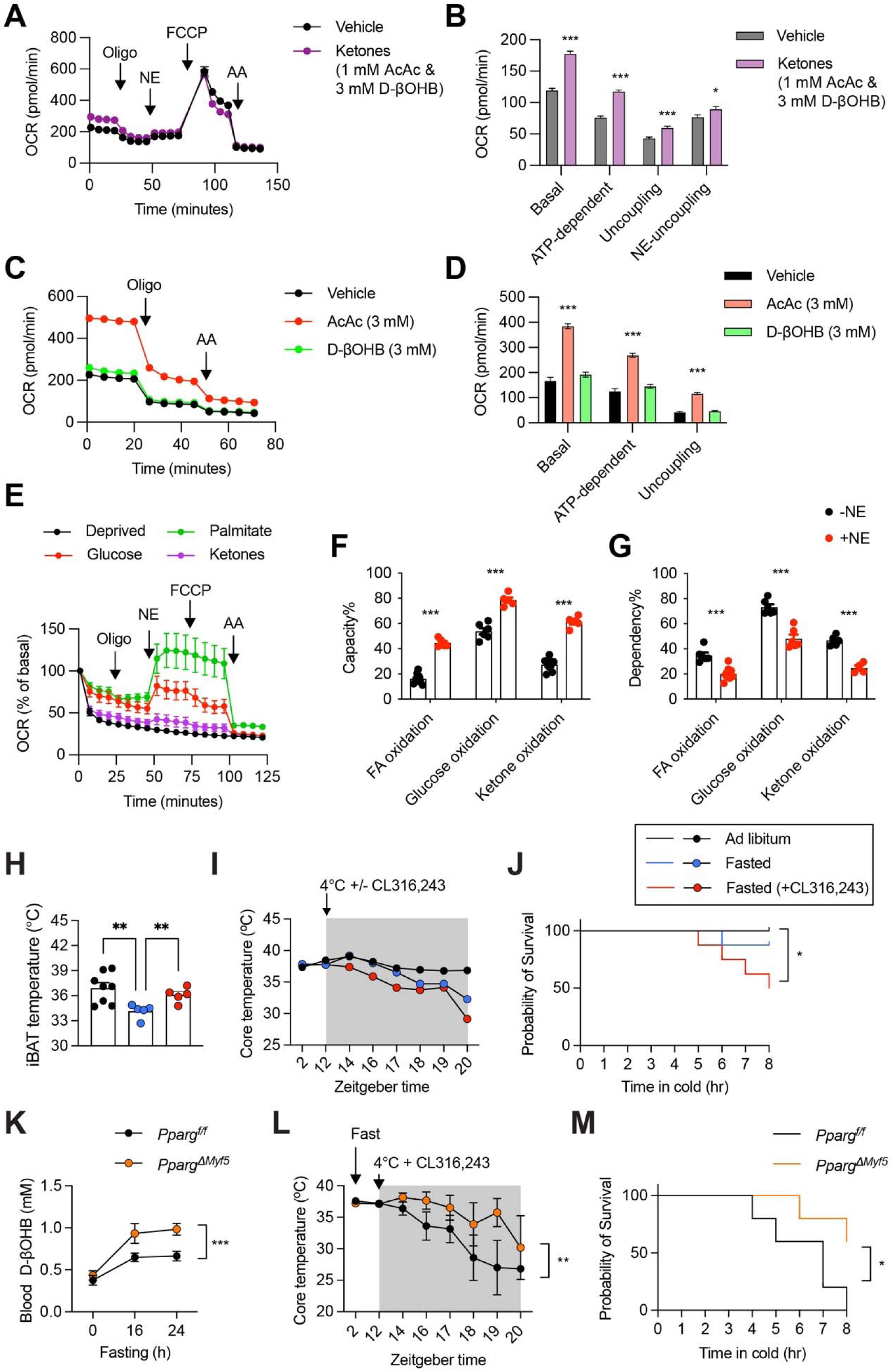
Ketone utilization by brown adipocytes for adaptive thermogenesis. (**A, B**) Primary brown adipocytes were pretreated with ketone bodies (3 mM D-βOHB, and 1 mM AcAc) in assay medium containing 5 mM glucose and 2 mM glutamine. Oxygen consumption rate (OCR) was measured at baseline and following oligomycin (5 µM), NE (2 µM), FCCP (7.5 µM), and Antimycin A (5 µM) injections (A). Basal mitochondrial, ATP-dependent, and uncoupled respiration were quantified (B). (**C, D**) Primary brown adipocytes were pretreated with individual ketone body (3 mM D-βOHB or AcAc), OCR was measured (C) and quantified for basal mitochondrial, ATP-dependent, and uncoupled respiration (D). (**E**) OCR measurement of primary brown adipocytes pretreated with glucose-deprived medium or supplemented with glucose (5 mM), mixed ketones (3 mM D-βOHB and 1 mM AcAc), or palmitate (0.2 mM) for 1h. (**F, G**) Metabolic flexibility in resting and NE (2 µM)-stimulated primary brown adipocytes that were pretreated with ketones-supplemented respiration assay media (3 mM D-βOHB,1 mM AcAc, 5 mM glucose, and 2 mM glutamine). (F) Substrate reserve capacity was determined for fatty acids (10 µM UK5099 and 500 nM pimozide), glucose (40 µM etomoxir and 500 nM pimozide), or ketenes (10 µM UK5099 and 40 µM etomoxir) when the oxidation of other two nutrients were inhibited. (G) Substrate oxidation dependence was determined for fatty acids, glucose, and ketones by the inhibition with etomoxir, UK5099, and pimozide, respectively. (**H-J**) C57BL/6J mice were AL fed or fasted at ZT2, exposure to cold at ZT12 with or without CL316,243 injection, and measured for iBAT temperature (H), rectal temperature (I), and probability of survival (J). (**K**) Blood D-βOHB levels of *Pparg^f/f^* and *Pparg*^Δ*Myf5*^ mice fasted overnight followed with 8 hour of cold challenge. (**L, M**) *Pparg^f/f^* and *Pparg*^Δ*Myf5*^ mice were fasted at ZT2, exposure to cold and injected with with at ZT12, and measured for rectal temperature (L), and probability of survival (M). Data are represented as mean ± SEM. * P < 0.05, ** P < 0.01, *** P < 0.001 by two-tailed unpaired Student’s t-test (B, F, G), one-way ANOVA (D, H), two-way ANOVA (I, K, L), and Kaplan-Meier Analysis (J, M). Please also see Figure S3.

Fasting suppresses thermogenic activity (Ruan *et al*., 2014), which can be an adaptive response for energy conservation and survival. To test this, we treated fasted and cold challenged mice with the β3-AR agonist CL316,243 in the dark cycle (**Fig. 3H**). Untreated mice generally could deal with the dual challenges by fast and cold; however, BAT overactivation by CL316,243 negatively affected body temperature control and the survival (**Fig. 3I, J**), supporting the notion that forced BAT overactivation during fasting causes an energy crisis in mice.

To validate that the detrimental effect of CL316,243 in the setting of fasting was mediated by brown adipocytes, we generated iBAT-atrophic *Pparg*^Δ*Myf5*^ mice by deleting the adipogenic master transcription factor PPARγ from the *Myf5*^+^ myogenic progenitors (Huang et al., 2023; Sanchez-Gurmaches and Guertin, 2014). *Pparg*^Δ*Myf5*^ mice showed higher levels of blood D-βOHB compared to *Pparg^f/f^* controls (**Fig. 3K**), supporting that BAT participates in systemic ketone disposal. *Pparg*^Δ*Myf5*^ mice did not possess iBAT to be activated by CL316,243, so they largely maintained body temperature and improved survival when challenged by fasting and cold in the dark cycle (**Fig. 3L, M**). Collectively, these results indicate that ketone utilization by BAT during fasting allows adaptative reduction in thermogenesis, which helps animals conserve energy for long-term survival in cold.

### Mitochondrial oxidation of AcAc in BAT

Exogenous D-βOHB has been shown to undergo both mitochondrial oxidization and incorporation into lipids in BAT (Agius and Williamson, 1981; Wright and Agius, 1983), which likely proceeds exclusively through an AcAc intermediate. To directly determine the metabolic fate of ketone bodies, we treated primary mouse brown adipocytes with 1 mM sodium AcAc or sodium [U-^13^C_4_]AcAc, in the absence or presence of NE, then employed ultra-high performance liquid chromatography (UHPLC) coupled to high mass accuracy, high mass resolution mass spectrometry (HRMS) to perform ^13^C stable isotope tracing. Measurements of the ^13^C enrichment of the TCA cycle intermediates found that citrate, αKG, succinate, fumarate, and malate were all ^13^C enriched 24.8 ± 5.6%, 25.9 ± 2.9%, 6.5 ± 0.9%, 13.8 ± 4.3%, and 14.7 ± 3.6%, respectively **(Fig. 4A)**. The most intense mass isotopologue detected in each metabolite was M+2, suggesting ketone bodies entered the TCA cycle through acetyl-CoA likely in a OXCT1-dependent manner, which is consistent with ketone bodies serving as an oxidative fuel in brown adipocytes. Interestingly, AcAc uptake, the fractional ^13^C enrichment, total pool size, and relative intensity of the M+2 isotopologue for each metabolite were all unchanged by NE treatment (**Fig. S4 and data not shown**), suggesting that ketone body oxidation in brown adipocytes likely proceeds in a NE-independent manner. Even so, NE can modulate the dependence and capacity of ketone-stimulated OCR (**Fig. 3F, G**), expectedly via its regulatory role on glucose and fatty acid oxidation (Ruan, 2020).

**Figure 4.**
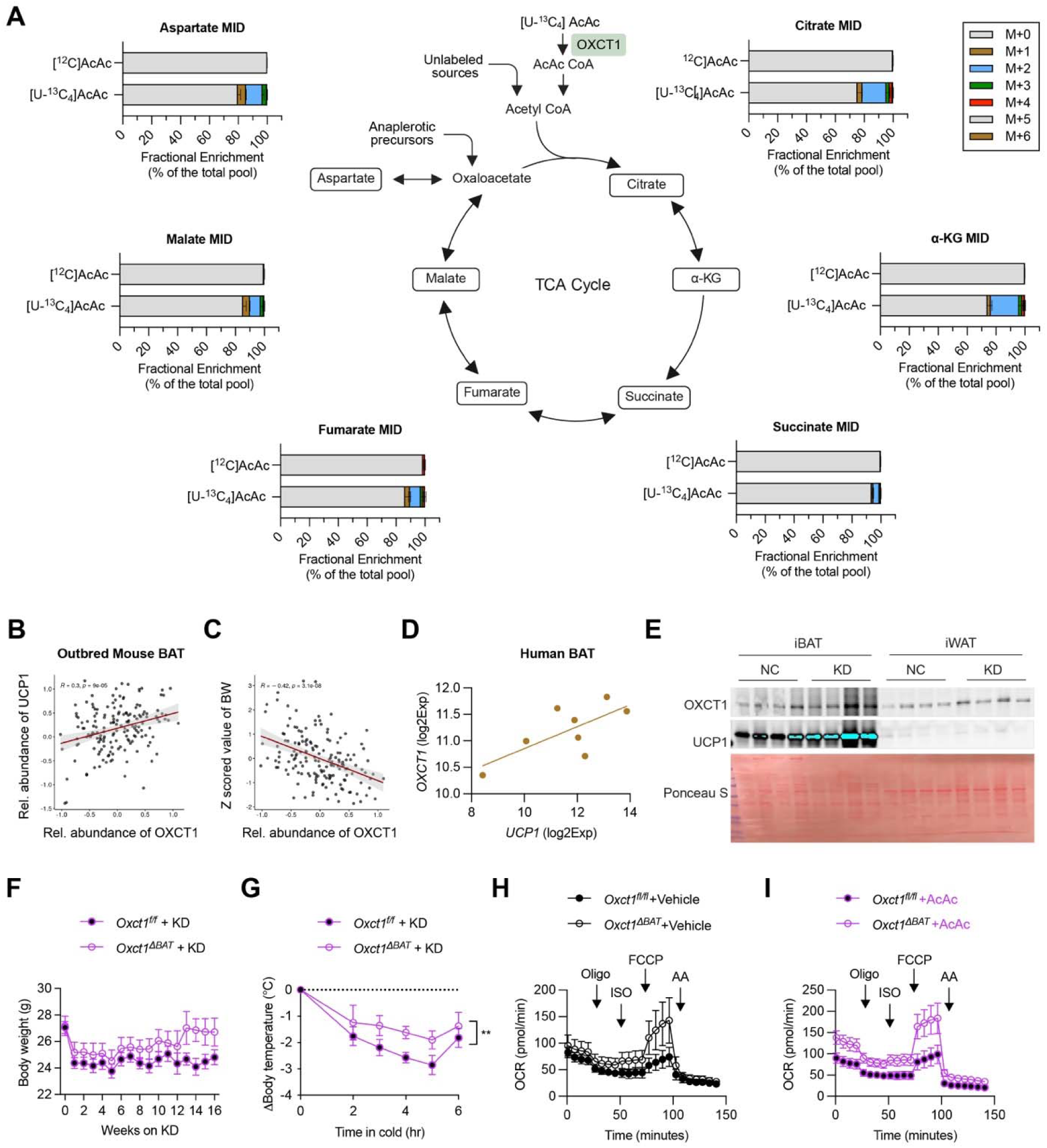
OXCT1-dependent ketone oxidation in brown adipocytes. (**A**) Mass isotopologue distribution (MID) of TCA cycle intermediates in primary brown adipocytes treated with 1 mM sodium [^12^C]AcAc or sodium [U-^13^C_4_]AcAc for 24 h. (**B, C**) Regression analysis of OXCT1 protein expression in BAT from a cohort of outbred mice with UCP1 (B) and body weight (C). (**D**) Correlation of OXCT1 and UCP1 protein expression in human BAT. (**E**) C57BL/6J mice were fed normal chow (NC) or ketogenic diet (KD). OXCT1 and UCP1 expression in interscapular BAT (iBAT) and inguinal WAT (iWAT) were determined by Western blotting. (**F, G**) *Oxct1^f/f^* and *Oxct1*^Δ*BAT*^ mice were fed with KD for 16 weeks. Body weight (F) and cold tolerance (G) were measured. (**H, I**) Seahorse assay of primary adipocytes differentiated from *Oxct1^f/f^* and *Oxct1*^Δ*BAT*^ BAT SVF cells in the absence (H) or presence (I) of 3 mM AcAc. Data are represented as mean ± SEM. * P < 0.05 by two-way ANOVA (G). Please also see Figure S4-6.

Mitochondrial OXCT1 converts AcAc to AcAc-CoA for subsequent oxidation. By analyzing BAT proteomics data from a cohort of genetically defined diversity outbred mice (Xiao et al., 2022), we found that OXCT1 positively correlates with UCP1 (**Fig. 4B**), UCP1’s co-operative partners, and key thermogenic activators (**Fig. S5**). On the other hand, OXCT1 expression in BAT from outbred mice was negatively correlated to body weight (**Fig. 4C**), fat mass, and the expression of positive driver of adiposity such as LEP and NPR3 (**Fig. S5**). In human BAT (Muller *et al*., 2016), OXCT1 showed strong correlative expression with UCP1 (**Fig. 4D**). To investigate if ketolysis in brown adipocytes is physiologically relevant, we generated BAT-specific *Oxct1*^Δ*BAT*^ knockout mice using the *Ucp1^Cre^* line (Kong et al., 2014). Western blotting of whole adipose tissue and isolated adipocytes confirmed OXCT1 deletion in BAT but not inguinal WAT (**Fig. S6A, B**). Fasting serum ketone levels was slightly higher in *Oxct1*^Δ*BAT*^ mice (**Fig. S6C, G**), implying that only a small portion of circulating ketones were oxidized in BAT mitochondria via OXCT1. We did not observe evident effect of OXCT1 deficiency on UCP1 expression, body weight and adipose tissue weight, or cold tolerance in both female and male mice fed with normal chow (**Fig. S6B-J**).

We postulated that ketone utilization by BAT is limited when other nutrients are abundantly and readily available. Inducing nutritional ketosis in mice by low-carbohydrate high-fat, ketogenic diet (KD) increased OXCT1 and UCP1 expression in BAT (**Fig. 4E**). We then fed control and *Oxct1*^Δ*BAT*^ mice with KD to induce ketosis and body weight loss (**Fig. 4F**). Notably, *Oxct1*^Δ*BAT*^ mice showed better cold tolerance when compared to controls (**Fig. 4G**), leading to the speculation that OXCT1-mediated oxidation may not be a favored route of ketone utilization in BAT. Consistent with this speculation, seahorse assay revealed an increase of basal, uncoupling, and reserve oxygen consumption in OXCT1-deficient brown adipocytes, especially when AcAc was provided exogenously (**Fig. 4H, I and Fig. S6K, L**). These results underscore a hypothesis that OXCT1-independent utilization of AcAc in BAT is driving thermogenesis. This shunt pathway is likely the diversion of ketones into lipogenesis, which remained hitherto unknown in BAT and was investigated next.

### AcAc supports fatty acid biosynthesis in BAT

We recently discovered in hepatocytes that ketone bodies contributed carbon directly to de novo lipogenesis (DNL) and fatty acid (FA) elongation, in a manner that was vital for maintaining polyunsaturated FA (PUFA) homeostasis in the liver (Queathem *et al*., 2025). Ketone body-sourced DNL and PUFA elongation is mediated by cytosolic acetoacetyl-CoA synthetase (AACS) that converts AcAc to AcAc-CoA (Bergstrom, 2023). Brown adipocytes had markedly higher expression of AACS than white adipocytes (**Fig. 5A**). In genetically defined diversity outbred mice (Xiao *et al*., 2022), AACS protein expression in BAT was positively correlates with the expression of UCP1 and its co-operative partners (**Fig. 5B and Fig. S5**), as well as ELOVL6 (**Fig. 5C**), an enzyme involved in the elongation of long-chain fatty acids and necessary for thermogenic capacity (Tan et al., 2015). Notably, AACS and OXCT1 expression is negatively correlated (**Fig. 5D**), suggesting the cytosolic lipogenesis and mitochondrial oxidation of ketone bodies could be exclusive events.

**Figure 5.**
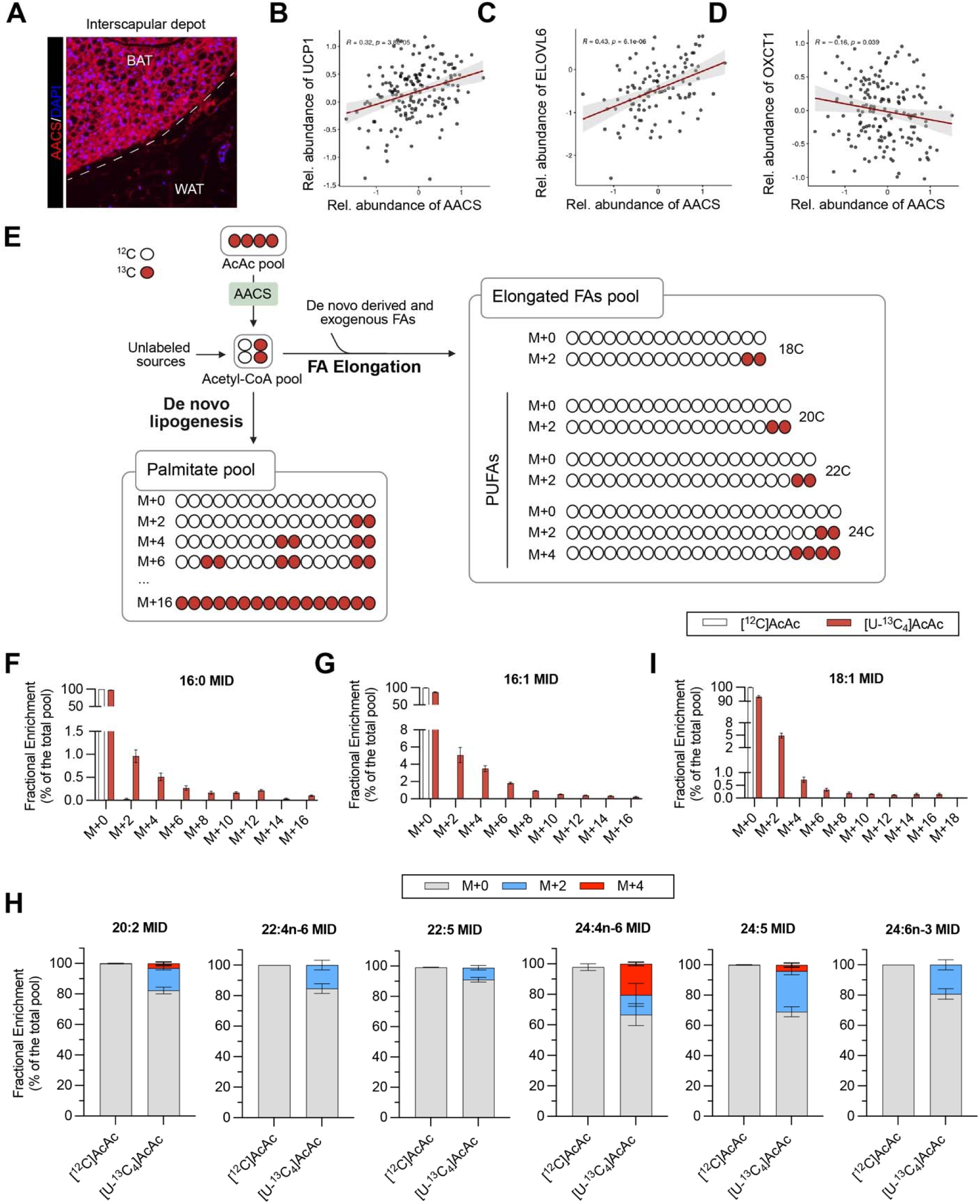
The lipogenic fate of AcAc in brown adipocytes. (**A**) Immunofluorescent staining of AACS in interscapular adipose tissue. BAT and nearby WAT were indicated. (**B-D**) Regression analysis of AACS protein expression in BAT from a cohort of outbred mice with UCP1 (B), ELOVL6 (C), and OXCT1 (D). (**E**) Lipogenic fate of [U-^13^C_4_] AcAc outlining the generation of [^13^C] Acetyl-CoA and subsequent incorporation into de novo lipogenesis and fatty acid elongation. (**F-I**) ^13^C-MID of de novo-synthesized (M+2, M+4, M+6,…,M+16) palmitate (F), palmitoleate (G), oleate (I), and elongated (M+2 or M+4) PUFAs (H) in primary brown adipocytes. Data are represented as mean ± SEM. Please also see Figure S7,8.

While prior reports employing ^14^C radioisotope tracers indicated that ketone bodies could contribute to BAT lipids (Agius and Williamson, 1981; Wright and Agius, 1983), the specific lipid species sourced by ketone bodies in brown adipocytes was not known. To study the lipogenic fate of ketone bodies in BAT metabolism, we returned to our ^13^C stable isotope tracing data and quantified the contribution of [U-^13^C_4_]AcAc to free FAs (FFAs) using reverse phase UHPLC-HRMS (**Fig. 5E**). We found that AcAc contributed to DNL in brown adipocytes, as evidenced by the ^13^C enrichment of the FFAs palmitate (16:0, 2.4 ± 0.5% of the total pool, **Fig. 5F**) and palmitoleate (16:1, 13.0 ± 2.4% of the total pool, **Fig. 5G**). We also detected ^13^C enrichment in numerous PUFA species including 20:2, 22:4, 22:5, 24:4, 24:5, and 24:6, ranging from 5-25% fractional ^13^C enrichment (**Fig. 5H**), demonstrating for the first time that ketone bodies contribute to FA elongation and PUFA biosynthesis in brown adipocytes. Similar to what was observed in primary hepatocytes, the ^13^C enrichment pattern of PUFAs from ketone bodies displayed primarily M+2 isotopologues (**Fig. 5H and Fig. S7**), reflecting entry of ^13^C carbon via elongases (ELOVLs), as opposed to the M+2, M+4, M+6… pattern introduced via fatty acid synthase (FAS) during DNL. Notably, we also detected ^13^C enrichment of monounsaturated FA (MUFA) oleate (18:1, 6.8 ± 1.3% of the total pool, **Fig. 5I**), which arise via a combination of DNL and FA elongation. While the characteristic M+2, M+4, M+6 pattern of DNL was encoded in oleate, the M+2 isotopologue was considerably more intense (4.9 ± 1.0% of the total pool), suggesting that while ketone bodies can contribute to both DNL and FA elongation, that ketones are selectively partitioned into elongation over DNL (**Fig. 5I**). we did not detect any effect of NE on the ^13^C enrichment of any FFA species (**Fig. S7**), indicating that both the oxidative and lipogenic fates of ketone bodies in brown adipocytes proceed via mechanisms that are independent of BAT-autonomous NE signaling.

To determine the systemic effect of lipid remodeling by BAT ketone metabolism, we performed metabolomics of arterial (A) and Sulzer’ venous (V) blood samples collected from fasted *Hmgcs2*^Δ*Hep*^ mice exposed to cold (**Fig. S8A**). Calculated log2 A/V ratios showed that oleate (18:1) reduced its net release from BAT (**Fig. S8B**). We also observed increased uptake of certain species of ceramides and lysophosphatidylcholine (LPC), specifically those containing MUFAs and PUFAs (**Fig. S8C-E**). These results point to a scenario that when endogenous ketone body-supported FA elongation and desaturation are absent, BAT consumes more unsaturated long-chain FAs (LCFAs) from the circulation to compensate for thermogenic function.

### Ketone driven lipid remodeling supports adaptive thermogenesis

Ketone bodies contribute to brown adipocyte lipogenic metabolism, likely via AACS. To examine AACS-mediated fatty acid biosynthesis in BAT thermogenesis, we obtained the whole-body *Aacs*^-/-^ knockout mice (**Fig. 6A**) and induced ketosis with ketogenic diet (**Fig. S9A, B**). Compared to controls, *Aacs*^-/-^ mice exhibited lower body temperature during cold exposure (**Fig. 6B**). To investigate BAT-specific role of AACS in thermogenesis, we differentiated brown adipocytes from BAT SVF cells for Seahorse respirometry. AACS deficiency attenuated basal, ATP-linked, and proton leak respiration (**Fig. 6C and Fig. S9C**), which was further exacerbated by the presence of AcAc (**Fig. 6D and Fig. S9D**). Similarly, freshly isolated BAT tissue from *Aacs*^-/-^ mice also demonstrated reduced OCR (**Fig. 6E**). Our results provide the evidence that lipid synthesis from ketones mediate by BAT-autonomous AACS is vital for oxygen consumption and thermogenesis.

**Figure 6.**
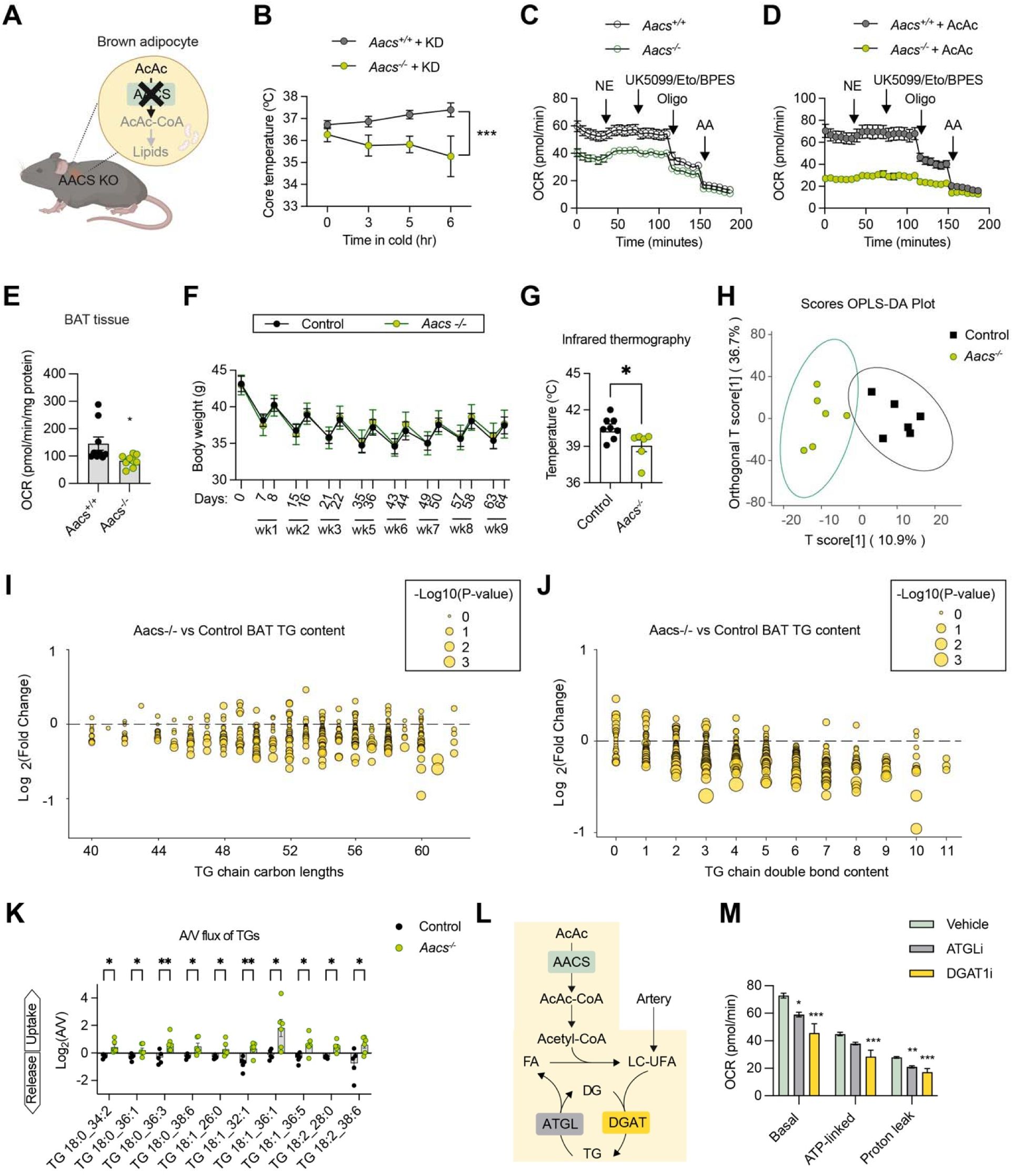
AACS-dependent lipid remodeling in BAT. (**A, B**) *Aacs*^-/-^ mice were obtained (A), fed with KD, and measured for cold tolerance (B). (**C, D**) Seahorse assay of primary brown adipocytes from *Aacs*^+/+^ and *Aacs*^-/-^ mice in assay medium containing 0 mM (C) or 3 mM (D) AcAc. (**E**) Seahorse assay of BAT explants from *Aacs*^+/+^ and *Aacs*^-/-^ mice. (**F, G**) *Aacs*^+/+^ and *Aacs*^-/-^ mice were first induced obesity with HFD and then subjected to ADF. (F) Body weight after AFD. (G) iBAT temperature measured by infrared camera. (**H-J**) Lipidomics was performed using BAT from HFD-ADF *Aacs*^+/+^ and *Aacs*^-/-^ mice collected on the fasting day. (H) OPLS-DA plot showing the variance of BAT lipid species. (I, J) Bubble plots showing changes of TGs with different carbon lengths (I) and double bounds (J). (**K**) A/V blood samples from HFD-ADF *Aacs*^+/+^ and *Aacs*^-/-^ mice on the fasting day were subjected for metabolomics. Log2A/V values were calculated to show the update (positive values) of TGs (**I**) Schematic view of AcAc-sourced, AACS-dependent Acetyl-CoA in driving FA modification and lipid cycling. (**M**) Seahorse measurement of basal, ATP-linked, and protein leak OCR in primary brown adipocytes treated with inhibitors to ATGL or DGAT. Data are represented as mean ± SEM. * P < 0.05, ** P < 0.01, *** P < 0.001 by two-tailed unpaired Student’s t-test (E, G, K), one-way ANOVA (M), or two-way ANOVA (B). Please also see Figure S9, 10.

Time-restricted eating (TRE) has recently emerged as an efficient and adherable treatment for obesity (Chaix et al., 2019; Chow et al., 2020); however, TRE renders mice more vulnerable to cold-induced hypothermia (van der Vinne *et al*., 2018; Zhang et al., 2020). We asked if ketone metabolism in BAT contributed to cold sensitivity in the setting of TRE. Subjecting high-fat diet-(HFD-) fed mice to alternate day fasting (ADF) prevented obesity (**Fig. S9E**), led to ketosis on fasting days (**Fig. S9F**), and upregulated AACS expression in BAT (**Fig. S9G, H**). ADF caused a similar reduction in body weight in HFD-fed control and *Aacs*^-/-^ mice (**Fig. 6F**). Yet, BAT from *Aacs*^-/-^ mice on the HFD-ADF treatment displayer lower temperature than that of control mice on the fasting day (**Fig. 6G**), supporting the role of AACS in BAT thermogenesis in mice upon TRE, independent of weight control.

To determine the consequence of AACS deficiency on lipid remodeling, we performed lipidomics of BAT tissues. Discriminant analysis by orthogonal partial least squares (OPLS-DA) uncovered systemic changes in the lipidome between control and *Aacs*^-/-^ BAT (**Fig. 6H**). Differential lipids predominantly consisted of TGs, ceramides, and glycerophospholipids (**Fig. S10A**). Analysis of lipid chain length imparted a clear downregulation of TGs consisting of LCFAs (**Fig. 6I**). Quantification of lipid chain unsaturation revealed that the more double bonds TGs contained, the less abundance they were in *Aacs*^-/-^ BAT (**Fig. 6J and Fig. S10B-D**). Furthermore, significantly differential lipids were enriched in pathways classified for thermogenesis, fat metabolism and insulin signaling (**Fig. S10E**). Taken together, ketone bodies prominently remodel the BAT lipid pool.

Due to the endogenous deficiency of PUFA synthesis from ketone bodies in *Aacs*^-/-^ BAT, it consumed more PUFA-TGs from the circulation, determined by metabolomics of AV samples (**Fig. 6K**). These results support a model that PUFAs derived from ketones bodies drive TG cycling and modification (**Fig. 6L**) (Wunderling et al., 2023). Indeed, the ability of mixed ketone bodies to promote brown adipocyte OCR was attenuated when TG synthesis and breakdown were suppressed by DGAT2 inhibitor and ATGL inhibitor, respectively (**Fig. 6M**). Collectively, these data indicate that the role of ketones in BAT metabolism and the thermogenic response may proceed via the modulation of PUFA pools. Prior research has stipulated the functional difference between DNL and FA elongation in BAT. While endogenous DNL strongly suppresses adipocyte thermogenesis (Guilherme et al., 2023; Schlein et al., 2021), the remodeling of fatty acid chain length is necessary for thermogenic recruitment of BAT (Tan *et al*., 2015; Westerberg et al., 2006). The preferential shunt of ketone-derived Acetyl-CoA to FA elongation and PUFA synthesis provides a mechanistic foundation to uncouple DNL and FA remodeling in BAT function and adaptation.

### BAT ketone metabolism mediates effects of time-restricted eating

Finally, we determined the relevance of ketone metabolism in human BAT. We isolated preadipocytes from human deep neck BAT and subcutaneous WAT and differentiated them into primary adipocytes (Arianti et al., 2024; Huang *et al*., 2023). Higher expression of BDH1, OXCT1, and AACS proteins could be observed in human brown adipocytes than white adipocytes (**Fig. 7A, B**). Similarly, the expression of *BDH1*, *OXCT1*, and *AACS* genes was abundant in adipocytes derived from deep neck BAT (**Fig. S11A**). Cyclic AMP (cAMP), mimicking adrenergic stimulation, did not significantly affect the levels of ketone metabolism genes and proteins (**Fig. 7A, B and Fig. S11A**), consistently with the tracing data showing minimal effects of NE on metabolic fates of ketone bodies. It suggests that the expression of ketone metabolism genes/proteins is prominently regulated by food availability, rather than cold-associated signals.

**Figure 7.**
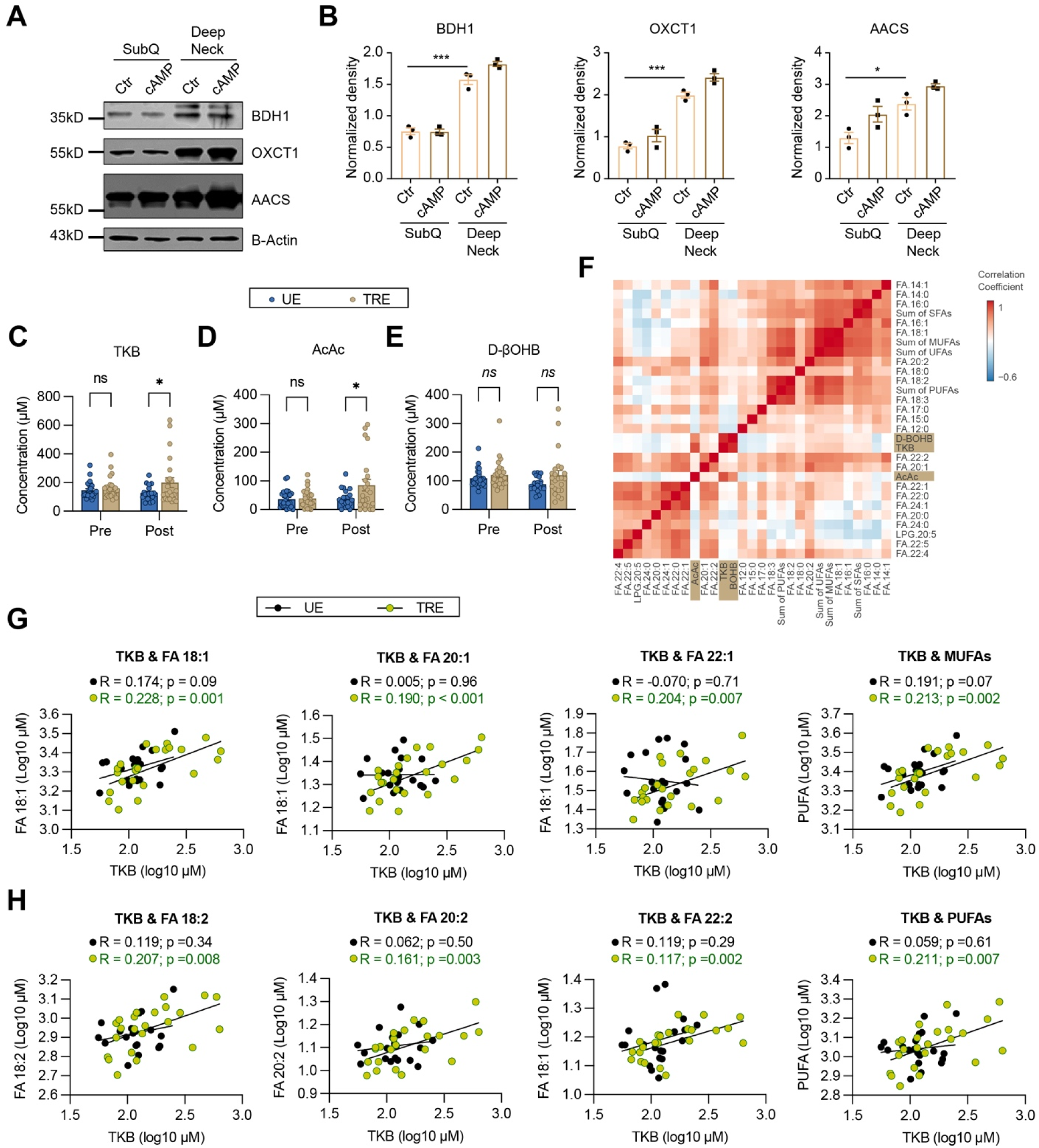
Association of plasma ketone bodies and unsaturated FAs in TRE humans. (**A, B**) Preadipocytes from human subcutaneous (SubQ) WAT and deep neck BAT were differentiated into adipocytes and treated with or without cAMP. BDH1, OXCT1, and AACS protein expression was determined by immunoblotting (A) and quantified for densitometry (B). (**C-E**) Measurement of TKB (C), AcAc (D), and D-βOHB (E) in plasma samples from patients with obesity and without diabetes who were subjected to unrestricted eating (UE) or time restricted eating (TRE). (**F**) Lipid species correlated with TKB, AcAc, and D-βOHB in post-treatment plasma samples from UE and TRE patients. (**G, H**) Regression analysis of post-treatment plasma AcAc level with selected and total MUFAs (G) and PUFAs (H) in UE and TRE patients. Data are represented as mean ± SEM. * P < 0.05, ** P < 0.01, *** P < 0.001 by two-way ANOVA (B-E). Please also see Figure S11.

While TRE and calorie restriction (CR) are both effective for weight reduction (Lin et al., 2023; Pavlou et al., 2023), TRE has a daily, prolonged fasting window that CR does not impose in humans. We speculated about the induction of ketone metabolism and lipid remodeling by TRE. To this end, we analyzed plasma samples obtained from a 12-week randomized-controlled trial (RCT: Clinicaltrials.gov NCT04259632) which assigned participants with obesity and without diabetes to either unrestricted eating (UE), CR or TRE (daily 8-hour eating window). (Oldenburg et al., 2025). For this study, we compared samples from the TRE (n=30) and UE (n=29) groups, pre- and post-intervention. All participants, regardless of intervention group, received a standardized evening meal to consume between 6-8 PM before their fasting blood draw (7-9 AM). At end-intervention, participants in the TRE group had higher fasting plasma levels of total ketone body (TKB) (**Fig. 7C**) and AcAc, but not D-βOHB (**Fig. 7D, E**). Neither ketone body levels or change in ketone body levels correlated with any observed changes in primary or secondary outcomes (data not shown). To further test the relationship between ketone bodies and lipids, we performed lipidomics of post-intervention samples and correlated these results with post-intervention TKB, AcAc, and D-βOHB levels (**Fig. S11B**). We found that ketone bodies predominantly clustered with FFAs (**Fig. 7F**). Importantly, TKB levels positively correlated with MUFAs (18:1, 20:1 and 22:1, **Fig 7G**) and PUFAs (18:2, 20:2 and 22:2, **Fig 7H**) only in the TRE group, but not the UE group. On the other hand, regression analysis found no correlation with saturated FAs (16:0, 18:0, and 20:0, **Fig. S11C**). While the extent to which these effects are mediated by thermogenic fat in humans remains unknown, our observations suggest that the availability of ketone bodies is associated with improvement in plasma lipid profiles in the setting of TRE, suggesting the potential contribution of ketones driving FA biosynthesis.

## Discussion

BAT is one of the most metabolically active and flexible organs in mammals. An extraordinary amount of macronutrients are used by BAT for thermogenic fuels and building blocks. The metabolic plasticity of BAT helped the establishment of endothermy and expansion of mammals into diverse environments (Levy and Leonard, 2022). However, the bioenergic response of BAT for thermogenic needs and its contribution to whole-body metabolism during fasting conditions lack comprehensive investigations (Reinisch *et al*., 2020). In this study, by integrating isotope tracing, metabolomics and genetic mouse models, we unveiled that BAT depends on ketene body utilization and lipogenic shunt to support thermogenic adaptation during fasting. BAT ketone metabolism not only confers the survival advantage for mice facing extreme environments but may also be associated with improvement in the circulating lipid profile in the setting of TRE.

Inter-organ communication is vital for metabolic physiology, BAT thermogenesis included. Fat mobilization from WAT can compensate BAT-intrinsic defects in lipogenesis or lipolysis (Chitraju et al., 2020; Schreiber et al., 2017; Shin et al., 2017). Importantly, FFAs liberated from WAT is essential to defend body temperature during fasting (Schreiber *et al*., 2017; Shin *et al*., 2017). During cold exposure, liver undergoes a metabolic switch to provide acylcarnitines for BAT thermogenesis (Simcox et al., 2017). Liver is also the primary site for glucose output and glycerol phosphorylation, which can fuel BAT mitochondrial respiration and lipid cycling, respectively (Chaves et al., 2012; Hankir and Klingenspor, 2018; Wang et al., 2020). The ketone body shuttle described in this study represents an elegant example of WAT-Liver-BAT axis in regulating thermogenesis. Fasting stimulates WAT lipolysis and the released FFAs are oxidized by liver. Hepatic ketogenesis then uses FA oxidation-derived Acetyl-CoA for the synthesis of AcAc, which is a key adaptive nutrient for BAT thermogenesis during fasting. Whether active mechanisms are engaged by the liver to prefer AcAc production over D-βOHB and by BAT to more efficiently use AcAc than D-βOHB requires future investigations. It is worth noting that BAT has detectable BDH1 expression and D-βOHB can be metabolized by BAT, including lipid incorporation (Agius and Williamson, 1981). In addition to being a metabolic fuel, ketone bodies, D-βOHB in particular, have moonlighting roles as signaling molecules linking to posttranslational modifications and epigenetic regulations (Ruan and Crawford, 2018). Our study has forged ahead for a better understanding of the multidimensional roles of ketone bodies in inter-organ crosstalk and BAT metabolism.

When it comes to ketone utilization, most would survey the mitochondrial oxidation via OXCT1. Using isotope of tracing in brown adipocytes, we provided the first evidence that AcAc incorporates to both TCA intermediates in the mitochondria and lipid biosynthesis in the cytosol. While the relative flux between the two compartments of ketone metabolism is still unclear, our functional interrogation using genetic mouse models suggests distinct thermogenic outcomes of the two fates of ketone metabolism. *Oxct1*^Δ*BAT*^ mice had improved body temperature control during nutritional ketosis and OXCT1-deficient brown adipocytes had higher OCR when supplemented with AcAc. Mechanisms by which OXCT1-mediated ketone oxidation suppresses thermogenesis remain unclear, but we suspect its potential interference with glucose and FA oxidation, as well as the competition with the lipogenic flux of ketones. On the other hand, results from *Aacs*^-/-^ mice and primary adipocytes demonstrate that AACS-dependent lipid biosynthesis from ketones is functional important for adaptative thermogenesis. Improved cold sensitivity observed in *Oxct1*^Δ*BAT*^ mice might be a result of increased lipogenic shunt when terminal oxidation was suppressed. It will be important in future experiments to investigate regulatory mechanisms for AACS expression, AACS activity, and the incorporation rate of ketone bodies into lipid in vivo.

AcAc contributes to DNL and FA elongation primarily via AACS-dependent formation of the cytosolic Acetyl-CoA pool. Fractional enrichment of [^13^C]AcAc was much higher in PUFAs than palmitate, suggesting that AcAc is preferentially sourced to FA modification than de novo synthesis. While the underlying mechanisms are still unknown, it is plausible that AACS interacts and cooperates with elongases (ELOVLs) for the Acetyl-CoA channeling. It warrants future research to examine the potential physical and functional interactions between AACS and ELOVLs in BAT. When hepatic ketogenesis or BAT ketone utilization were interrupted in mice, profound perturbations in PUFA-containing TG and glycerophospholipid pools occurred in BAT. Our results indicate that ketone-sourced PUFAs can enter the Lands cycle and the TG-FA cycle to remodel phospholipids and TG. Such remodeling is essential for BAT function, because it is involved in mitochondrial function, energy expenditure, and the synthesis of lipokines (Lynes et al., 2019; Murphy and Folco, 2019; Sharma et al., 2024).

In summary, our study establishes a ketone shuttle between liver and BAT that is functionally indispensable for animals coping with nutrient deficit and cold stress. This liver-BAT axis serves as both a survival mechanism and a potential therapeutic target to improve lipid profiles and metabolic flexibility.

## Acknowledgements

We thank the Physiology Core of University of Minnesota for services related to controlled temperature room, metabolic cages and EchoMRI analyses, CAM-SU of Soochow University for generating genetic knockout mouse models and technical support in mouse phenotyping, and Dr. Ferenc Győry for the surgical removal of human BAT and WAT biopsies. This work was supported by American Heart Association (AHA) Innovative Project Award (24IPA1270908), National Institutes of Health (NIH, R01 AI162791), and Department of Integrative Biology & Physiology Grant Accelerator Program to H.-B.R, NIH (R01 AG069781 and R01 DK091538) to P.A.C. Z.M. was supported by AHA predoctoral fellowship (24PRE1196372). E.D.Q. was supported by National Heart, Lung, and Blood Institute training grant (T32 HL166142). E.K. and R.A. was supported by the National Research, Development and Innovation Office (NKFIH-FK145866 and PD146202) of Hungary. The clinical trial comparing TRE to UE was supported by the UMN CTSA Award (UM1TR004405-01A1) from the National Center for Advancing Translational Sciences, the NIH (R01DK124484) to L.S.C. L.S.C. and D.G.M received support from the Allen Foundation. A.B was supported by 2024 Research Infrastructure Investment Program from University of Minnesota.

## EXPERIMENTAL MODEL AND STUDY PARTICIPANT DETAILS

### Animals

All animal experiments were approved by the institutional animal care and use committee of the University of Minnesota and Soochow University. Animals were housed in a 14h light/10hdark cycle (6am-8pm light) and temperature- (20-22°C) controlled room with free access to water and normal chow (Teklad #2018) unless otherwise indicated.

*Hmgcs2*^Δ*Hep*^ mice were generated by injecting adeno-associated virus (AAV) with thyroid hormone-binding globulin (TBG) promotor driven iCre (1.5×10^11^ genomic copies in 0.1 ml volume, Vector Biolabs) into *Hmgcs2^f/f^* mice through tail vein. Control mice were generated by injecting AAV-TBG-eGFP into *Hmgcs2^f/f^*mice.

*Bdh1* global knockout mice (*Bdh1-Tm1a*, Knockout First) were made through injecting mutant mouse ES cells into mouse blastocysts. The positive insertion chimeric mice were then bred with wild type C57BL/6N mice to obtain *Bdh1-Tm1a* mice (abbreviated as *Bdh1^-/-^*). To generate Liver-specific BDH1 KO mice, *Bdh1^+/-^* mice were first crossed with *Flper* mice and littermates with then bred with wild type C57BL/6N mice to obtain *Bdh1-Tm1c* mice (abbreviated as *Bdh1^f/f^*). Liver-specific BDH1 KO mice were generated by crossing *Bdh1^f/f^* mice with Albumin (*Alb*)*-Cre* mice. Muscle-specific BDH1 KO mice were generated by crossing *Bdh1^f/f^* mice with muscle creatine kinase (*MCK*)*-Cre* transgenic mice.

*Pparg*^Δ*Myf5*^ mice were generated by crossing *Pparg^f/f^* mice (Jackson Laboratory # 004584) with *Myf5-Cre* mice (Jackson Laboratory # 007893). *Oxct1*^Δ*BAT*^ mice were generated by crossing *Oxct1^f/f^* mice with *Ucp1-Cre* mice (Jackson Laboratory # 024670). *Aacs*^-/-^ mice were sourced from the Mutant Mouse Resource & Research Centers (MMRRC, Cat# 042183-JAX).

### Housing Temperature Modifications

All housing temperature modifications, including cold challenge or thermoneutral conditions, were conducted in a walk-in environmental room at the IBP Phenotyping Core Facility, University of Minnesota.

### Alternate day fasting regimen

Mice were initially fed a 60% HFD (Research Diets, D12492) to induce obesity, with metabolic markers monitored throughout. Following that, animals underwent an alternate day fasting (ADF) regimen, remaining on HFD. During ADF, mice were transferred daily at ZT10 (4 hours before lights off) between cages with or without food, with continuous access to water.

### Human BAT

Tissue collection was approved by the Medical Research Council of Hungary (20571-2/2017/EKU) followed by the EU Member States’ Directive 2004/23/EC on presumed consent practice for tissue collection. All experiments were carried out in accordance with the guidelines of the Helsinki Declaration. All participants signed an informed consent before the surgical procedure. During the thyroidectomy, a pair of DN and SubQ adipose tissue samples was obtained to rule out inter-individual variations. Patients with known diabetes, malignant tumor, or with abnormal thyroid hormone levels at the time of surgery were excluded.

Human adipose-derived stromal cells (hASCs) were isolated from SubQ and DN fat biopsies as described previously and were differentiated into matured adipocytes following a protocol applying insulin, cortisol, T3, dexamethasone, and short-term rosiglitazone treatment (Arianti *et al*., 2024; Toth et al., 2020). After 14 days of differentiation, adipocytes were treated with a single bolus of 500 µM db-cAMP (D0260) for 10 h to mimic *in vivo* cold-induced thermogenesis. Global gene expression pattern of the differentiated and activated adipocytes was analyzed by bulk RNA-sequencing as previously described (Arianti *et al*., 2024).

### Human subjects

We had previously conducted a 12-week randomized controlled trial (Clinicaltrials.gov NCT04259632, Oldenburg et al., 2025) comparing three dietary interventions: 8-hour Time-Restricted Eating (TRE), 15% Caloric Restriction and Unrestricted Eating (UE). Eighty-eight participants (mean age 43.2±10.5 years, BMI 36.2±5.1 kg/m², 54.5% female, 84.1% white, baseline eating window of 14.4± 1.7 hours) were enrolled, with 81 (92%) completing the study. Participants were randomized to one of three arms: TRE with a self-selected daily 8-hour eating window, CR with a 15% reduction from baseline caloric intake, or UE serving as control. The actual eating windows achieved were 9.8 hours [95% CI: 9.0-10.6] for TRE, 12.9 hours [11.9-13.9] for CR, and 11.8 hours [11.0-12.7] for UE.

For the purpose of this study, we restricted our analysis to comparing the TRE (n=30) vs UE (n=29) groups. Plasma ketones were measured in the fasting state at baseline and end-intervention. Regardless of intervention group, all participants received a standardized evening meal with instructions on consuming the meal at 6-8 pm prior to the fasting blood draw (generally between 7-9 AM). The primary outcome was change in body weight. As reported in the primary paper (Oldenburg *et al*., 2025), after the 12 week intervention, weight changes compared to the UE group (n=29) were -1.4 kg [95% CI: -4.5 to 1.7; p=0.53] for TRE (n=30). Secondary outcomes included body composition measured by dual-energy X-ray absorptiometry (DXA), caloric intake assessed through 24-hour dietary recall, and glycemic measures including HbA1c, insulin sensitivity determined by hyperinsulinemic-euglycemic clamp, and glucose patterns via continuous glucose monitoring.

## METHOD DETAILS

### Infrared thermography and core temperature

Interscapular skin temperature was measured by anesthetizing mice with isoflurane and quickly capturing images using a FLIR C2 thermal camera. The average skin temperature within the interscapular region was analyzed using FLIR Thermal Studio. Core temperature was measured using a rectal probe (Physitemp), with recordings taken at the specified time points.

### Body composition and energy expenditure

Mice body composition was measured using Echo MRI (Echo Medical Systems) immediately before their introduction to indirect calorimetry cages for energy expenditure assessment (Oxymax, Columbus Instruments). Mice were housed in the calorimetry cages for consecutive days at 22LJ±LJ1°C or 4LJ±LJ1°C, as specified in the experimental design. Data collected during the initial acclimation period was excluded from the analysis. For ADF (alternate-day fasting) experiments, food was added or removed from the calorimetry cages at 4:00 pm (ZT0). Raw data were analyzed using the CalR.

### Arteriovenous (AV) blood sampling and processing

Arteriovenous (AV) blood collection was performed as previously described, with slight modifications. Briefly, mice were anesthetized with isoflurane, and venous blood was collected from Sulzer’s vein using 100 µL microvettes (Braintree Scientific) followed by arterial blood collection from the left ventricle using 27G 1 mL syringes. Blood samples were left undisturbed on ice for 20 minutes (in microvettes and microcentrifuge tubes, respectively) before centrifugation at 2,000g for 10 minutes at 4°C. The resulting serum samples were stored at - 80°C and sent to the Center for Metabolomics and Proteomics at the University of Minnesota for quantitative MS-based metabolomics analysis using the Biocrates platform (MxP Quant 500 XL). Biocrates outputs were uploaded into MetaboAnalyst for further analysis. For a subset of samples, ketone bodies (cyclic enzymatic assay, Wako Autokit Total Ketone Bodies and 3-HB), glucose, and triglycerides (both colorimetric assays) were quantified using commercially available micro assay kits. Collected interscapular brown adipose tissue samples were subjected to quantitative lipidomics profiling using ultra-performance liquid chromatography-tandem mass spectrometry (UPLC-MS/MS) platform (Metware Biotechnology Inc.).

### Mouse stromal vascular fraction (SVF) isolation, culture, and differentiation

SVF cells were isolated from the iBAT of 6–8-week-old mice as described previously with modifications (Oeckl et al., 2020). iBAT was carefully harvested to minimize erythrocyte contamination and rinsed in antibiotic-containing PBS (40 µg/mL penicillin-streptomycin, 40 µg/mL gentamicin, and 500 ng/mL amphotericin B). Up to three iBAT samples were minced thoroughly with sterile scissors in a small amount of digestion buffer inside a round-bottom microcentrifuge tube. The minced tissue was transferred to a 15 mL tube containing 7 mL of pre-warmed digestion buffer (DMEM/F12 with 1 mg/mL collagenase I, 1% FBS, 1% HEPES, and 1% penicillin-streptomycin). Digestion was performed at 37°C in an orbital shaker at 180 rpm for up to 45 minutes, or until no visible tissue fragments remained. The digested iBAT was filtered through a 100 µm mesh, and the remaining content in the tube was rinsed with 7 mL of wash buffer (HBSS containing 3.5% (w/v) BSA, 40 µg/mL penicillin-streptomycin, 40 µg/mL gentamicin, and 500 ng/mL amphotericin B). The solution was centrifuged twice at 300g for 5 minutes at room temperature, with careful inversion of the tube between centrifugations. The supernatant was discarded, and the pellet was resuspended in 2 mL of ACK lysis buffer for 3–5 minutes to lyse erythrocytes. Next, 13 mL of wash buffer was added to the resuspended pellet, followed by centrifugation at 500g for 5 minutes at room temperature. The cell pellet was resuspended in growth media (DMEM-high glucose supplemented with 20% (v/v) FBS, 40 µg/mL penicillin-streptomycin, 40 µg/mL gentamicin, and 500 ng/mL amphotericin B), filtered through a 70 µm mesh, and seeded into a 10 cm petri dish. Growth media was replaced the next day and subsequently every other day until the cells reached ∼80% confluency (∼4–5 days for iBAT from ∼3 mice). To induce differentiation, cells were treated with induction media (DMEM-high glucose with 10% (v/v) FBS, 40 µg/mL penicillin-streptomycin, 40 µg/mL gentamicin, 850 nM insulin, 1 nM T3, 1 µM rosiglitazone, 1 µM dexamethasone, 500 µM IBMX, and 12 µM indomethacin) for 2 days. To minimize well-to-well variability in downstream assays (e.g., Seahorse assay, uptake assay), cells were initially differentiated in the original petri dish. On day 2 of differentiation, cells were detached using 0.25% trypsin-EDTA and re-plated into micro-well plates at the desired density. Differentiation was continued until day 7–9, with the media being replaced every other day.

### Acetoacetate synthesis

Unlabeled and [U-^13^C_4_]acetoacetate was synthesized from unlabeled ethylacetoacetate (Sigma-Aldrich, W241512) or [1,2,3,4-^13^C_4_]ethylacetoacetate (CIL, CLM-3297-PK) via base-catalyzed hydrolysis as described previously (Puchalska et al., 2019; Queathem *et al*., 2025). Briefly, 8 mL of 1M sodium hydroxide was added slowly to 1 mL of ethyl acetoacetate with continuous stirring on a hotplate. After 30 minutes at 60°C, the reaction mixture was cooled on ice, and the pH was promptly adjusted to 8 using 36.5-38% HCl. The synthesized acetoacetate was then filter-sterilized and aliquoted for storage at -80°C. The concentration of acetoacetate was measured using a cyclic enzymatic reaction with the Wako Autokit Total Ketone Bodies.

### AcAc consumption assay

Primary brown adipocytes were isolated and cultured as described above. Cells were treated with 1 mM sodium acetoacetate (AcAc) in the absence and presence of 2 uM NE for 24 hours, then conditioned media was collected and total ketone bodies were quantified via UHPLC-MS/MS as previously described (Queathem *et al*., 2025).

### Metabolite extraction and sample preparation

Metabolite extraction and sample preparation were performed as previously described (Queathem *et al*., 2025). Briefly, conditioned media was aspirated from cultured cells, then cells were washed twice with 1mL warmed (37°C) PBS (without MgCl_2_ or CaCl_2_), twice with warmed (37°C) cell culture grade H_2_O, then culture dishes were flash frozen in liquid nitrogen to quench metabolism. Cells were harvested in 500 μL of ice cold (-20°C) LCMS grade MeOH then dried to completion in a SpeedVac at room temperature. Metabolites were extracted into 1 mL of ice cold (-20°C) LCMS grade MeOH:ACN:H_2_O (2:2:1), dried to completion in a SpeedVac, then were resuspended in 40 μL of LCMS grade ACN:H_2_O (1:1) and transferred to a LCMS glass vial for analysis. For free fatty acid (FFA) analysis, 10 uL of extracted metabolites were dried down under N_2_ (g), then FFAs were derivatized with N-[4-(Aminomethyl)phenyl]pyridinium (AMPP) as previously described (Queathem *et al*., 2025; Wang et al., 2013). FFA-AMPP derivatives were resuspended in 100 µL CHCl_3_:MeOH (1:1) and transferred to a LCMS glass vial for analysis.

### 13C stable isotope tracing analysis

Extracted metabolites and lipids were separated using a Thermo Fisher Scientific Vanquish UHPLC system and were detected using full-scan high mass accuracy high mass resolution mass spectrometry (HR-MS) on a Thermo Fisher Scientific QExactive plus hybrid quadrupole-orbitrap mass spectrometer, fitted with a heated electrospray ionization source. Polar metabolites were separated on an Atlantis Premier BEH Z-HILIC column (2.1 x 100 mm, 1.7 µm) (Waters, 186009979) and detected in negative ionization mode. Mobile phase A (MPA) was 15 mM ammonium bicarbonate in 100% H_2_O, pH 9. MPB was 15 mM ammonium bicarbonate in 90% ACN, 10% H_2_O, pH 9. Total run time was 10 min, flow rate was 0.5 mL/min for the first 6 minutes then was increased to 1 mL/min for the remainder of the run. Column chamber was 30°C. Mobile phase gradient was as follows: 0 to 0.1 min, 90% MPB; 0.1 to 5 min, 90 → 65% MPB; 5 to 6 min, 65% MPB; 6 to 6.5 min, 65 → 90% MPB; 6.5 to 10 min, 90% MPB. FFA-AMPP derivatives were separated on a Cortecs UPLC T3 C18 silica column (2.1 x 150 mm, 1.6 μm) (Waters, 186008500) and detected in positive ionization mode. MPA was 100% H_2_O, 0.1% formic acid. MPB was 50% IPA, 50% ACN, and 0.1% formic acid. The total run time was 24 min, flow rate was 0.4 mL/min for the first 8 min and then was decreased to 0.1 mL/min for the remainder of the run. Column chamber was 40°C. Mobile phase gradient was as follows: 0 to 0.1 min, 2% MPB; 0.1 to 4 min, 2→98% MPB; 4 to 7 min 98% MPB; 7 to 8 min, 98→2% MPB; 8 to 8.5 min, 2% MPB; 8.5 to 9.5 min, 2→98% MPB; 9.5 to 12.5 min, 98% MPB; 12.5 to 13.5 min, 98→2% MPB; 13.5 to 14 min, 2% MPB; 14 to 15 min, 2→98% MPB; 15 to 18 min, 98% MPB; 18 to 19 min, 98→2% MPB; and 19 to 24 min, 2% MPB. For all analyses, both blanks and pooled quality control (QC) samples were injected periodically throughout the run. Blanks were ACN:H_2_O (1:1) for polar metabolite analysis, or CHCl_3_:MeOH (1:1) for FFA-AMPP analysis. The QC sample was a pooled sample of all unlabeled and ^13^C-labeled samples. For analyte identification, a mix of authentic standards for expected analytes was injected, as well as a pooled sample consisting only of unlabeled samples, which was analyzed via data-dependent analysis MS/MS as previously described (Queathem *et al*., 2025). To determine isotopic enrichment, ^13^C mass isotopologues were separated via HR-MS, then the ^13^C isotopic envelope was profiled based on the diagnostic shift in m/z (Δm/z = 1.0033 Da). Raw ion counts were plotted against retention time to generate extracted ion chromatograms then the area under the curve was extracted, summed, and expressed as a percentage of the total pool. For polar metabolites, Thermo Compound Discoverer 3.3 was used. For FFA-AMPP derivatives, each isotopologue was manually integrated using Thermo Xcalibur Quan Browser. Analyte identification was based upon predicted m/z, retention time compared to authentic external standards, and MS/MS fragmentation pattern compared to standards or online databases. ^13^C enrichment was determined for putatively identified features after natural abundance correction. Fractional intensities of each mass isotopologue were then graphed as a function of ^13^C content to generate mass isotopologue distributions (MIDs) or summed together to calculate fractional ^13^C enrichment of the total pool.

### Seahorse respirometry

Oxygen consumption rate (OCR) in differentiated brown adipocytes was measured using a Seahorse XFe96 Analyzer (Agilent). One hour before respirometry, differentiation media was replaced with respiration assay media (DMEM D5030, supplemented with 5 mM glucose, 31 mM NaCl, 2 mM GlutaMax™, 5 mM HEPES, and 2% (w/v) fatty acid-free BSA). Ketone bodies, palmitate, and inhibitors were added to the respiration assay media as outlined in each experiment (AcAc 3 mM, βOHB 3 mM, combined AcAc and βOHB in a 1:3 ratio at 3 mM, palmitate 1mM / 1.4% BSA, Atglistatin 40 µM, Triacsin C 5 µM, PF-064224439 40 µM, and 2-DG 50 mM). Cells were incubated at 37°C in a non-CO2 incubator. A combination of stimulatory and inhibitory agents was injected through the injection ports to assess ATP-linked respiration, proton leak, and maximal respiration (oligomycin 5 µM, isoproterenol 0.5 µM, norepinephrine 2 µM, FCCP 7.5 µM, antimycin A 5 µM, and rotenone 5 µM).

Ex vivo respirometry of iBAT biopsies was performed as previously described (Mackert et al., 2022) using Seahorse XFe96 Analyzer (Agilent). Micro restrainers were 3D printed at the University of Minnesota Imaging Center using Stereolithography 3D printing technology.

### Real-time RT-PCR

Frozen tissues and cultured cells were lysed with Trizol and reverse transcribed into cDNA with the iScriptTM cDNA Synthesis Kit. Real-time RT-PCR was performed using iTaqTM Universal SYBR Green Supermix and gene-specific primers on a Bio-Rad C1000 Thermal Cycler.

### Western blot

Frozen tissues and cultured cells were lysed with 1X RIPA lysis buffer with protease inhibitor, boiled with 2X Laemmli buffer, and subjected to Western blotting (20-50µg protein sample).

## QUANTIFICATION AND STATISTICAL ANALYSIS

Results are shown as mean ± SEM. N values and statistical analysis methods are described in figure legends. The statistical comparisons were carried out using two-tailed unpaired Student’s t test and one-way or two-way ANOVA with indicated post hoc tests with GraphPad Prism 10. Differences were considered significant when p < 0.05. ∗, p < 0.05; ∗∗, p < 0.01; ∗∗∗, p < 0.001.

## Notes

### Competing Interest Statement

The authors have declared no competing interest.

