## Supplemental Figures for "Ketone body driven lipid remodeling supports thermogenic adaptation to fasting"

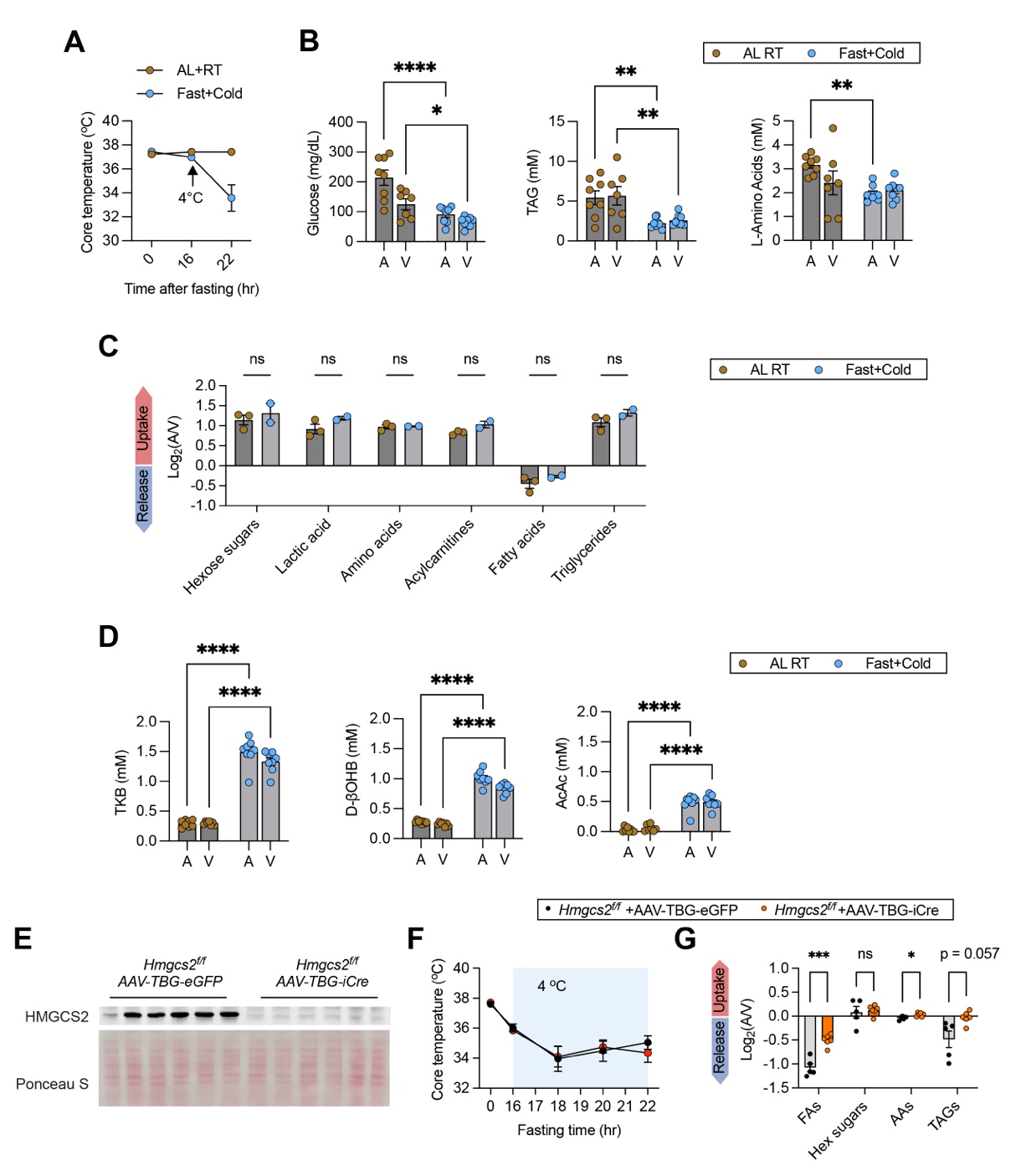

**Figure S1. Related to Figure 1. Hepatic ketogenesis and BAT thermogenic adaptation.**

(**A**) Mice were either AL fed or fasted overnight for 16h and exposed to RT or 4^o^C cold for 8h, respectively. Core body temperature was measured using a rectal probe (n = 8).

(**B**) Glucose, TG, and L-amino acids levels in left ventricle (A) and Sulzer’s vein (V) blood samples from AL-RT and Fast-Cold mice were measured with commercial kits.

(**C**) Log2 values of metabolites in A/V samples AL-RT and Fast-Cold mice, determined by metabolomics.

(**D**) TKB, D-βOHB, and AcAc levels in A/V samples from AL-RT and Fast-Cold mice.

(**E**) Immunoblotting validating knockdown of HMGCS2 in AAV-TBG-iCre injected *Hmgcs^f/p^* mice.

(**F, G**) Overnight fasted *Hmgcs2^ΔHep^* mice were cold exposed in the morning for 8 h. (F) Rectal temperature. (G) Log2 values of metabolites in A/V samples determined by metabolomics.

Data are represented as mean ± SEM. * P < 0.05, ** P < 0.01, *** P < 0.001, **** P < 0.0001 by two-tailed unpaired Student’s t-test (G) and two-way ANOVA (B, D) and n.s., not significant.

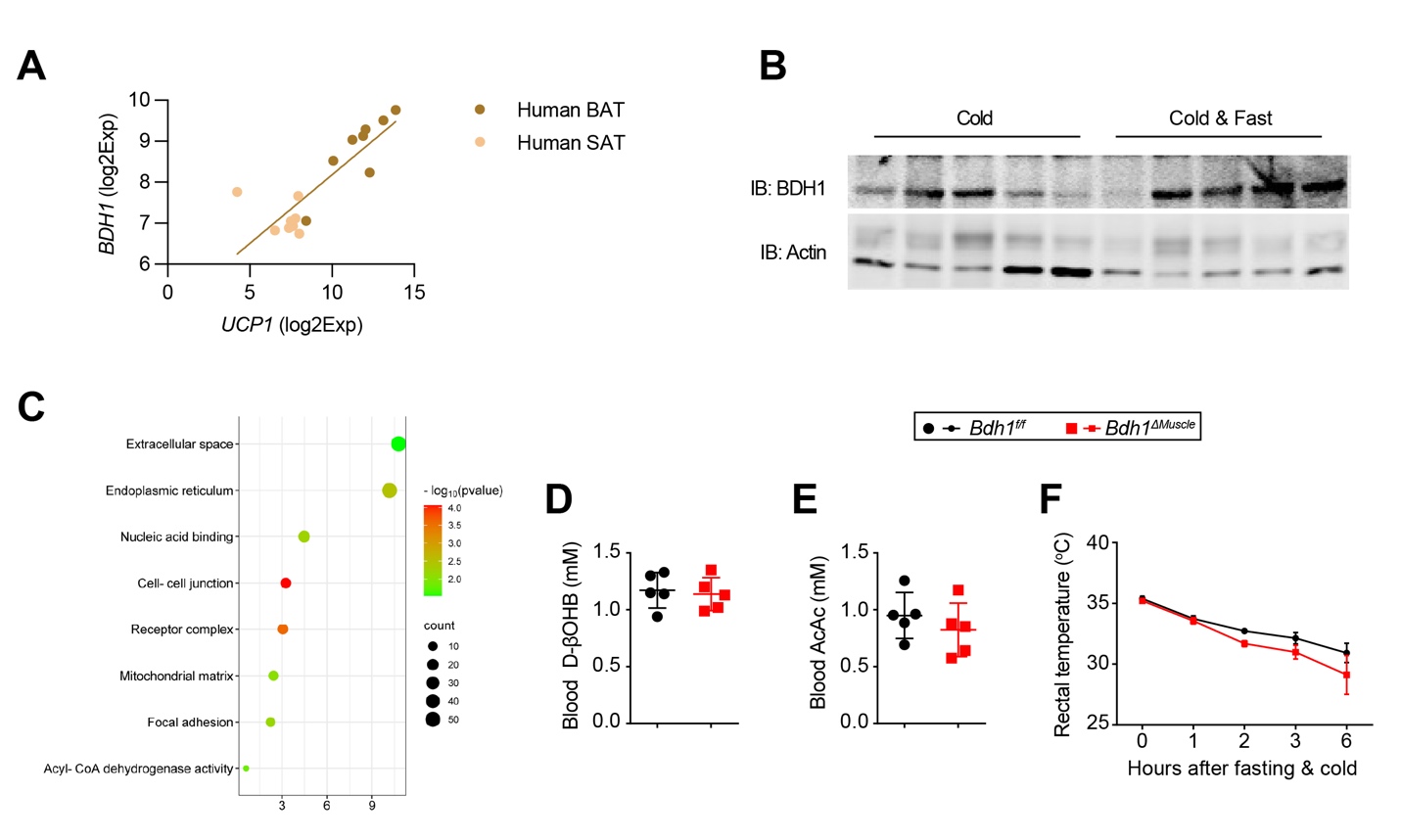

**Figure S2. Related to Figure 2. Role of BDH1.**

(**A**) Correlation of BDH1 and UCP1 protein expression in human BAT and subcutaneous adipose tissue (SAT).

(**B**) Expression of BDH1 in BAT from mice overnight challenged with cold or cold together with fasting.

(**C**) Pathway analysis of differentially expressed genes in BAT from *Bdh1^f/f^* and *Bdh1^ΔHep^* mice.

(**D-F**) Blood D-βOHB (D), blood AcAc (E), and cold tolerance (F) in *Bdh1^f/f^* and *Bdh1^ΔMuscle^* mice.

Data are represented as mean ± SEM.

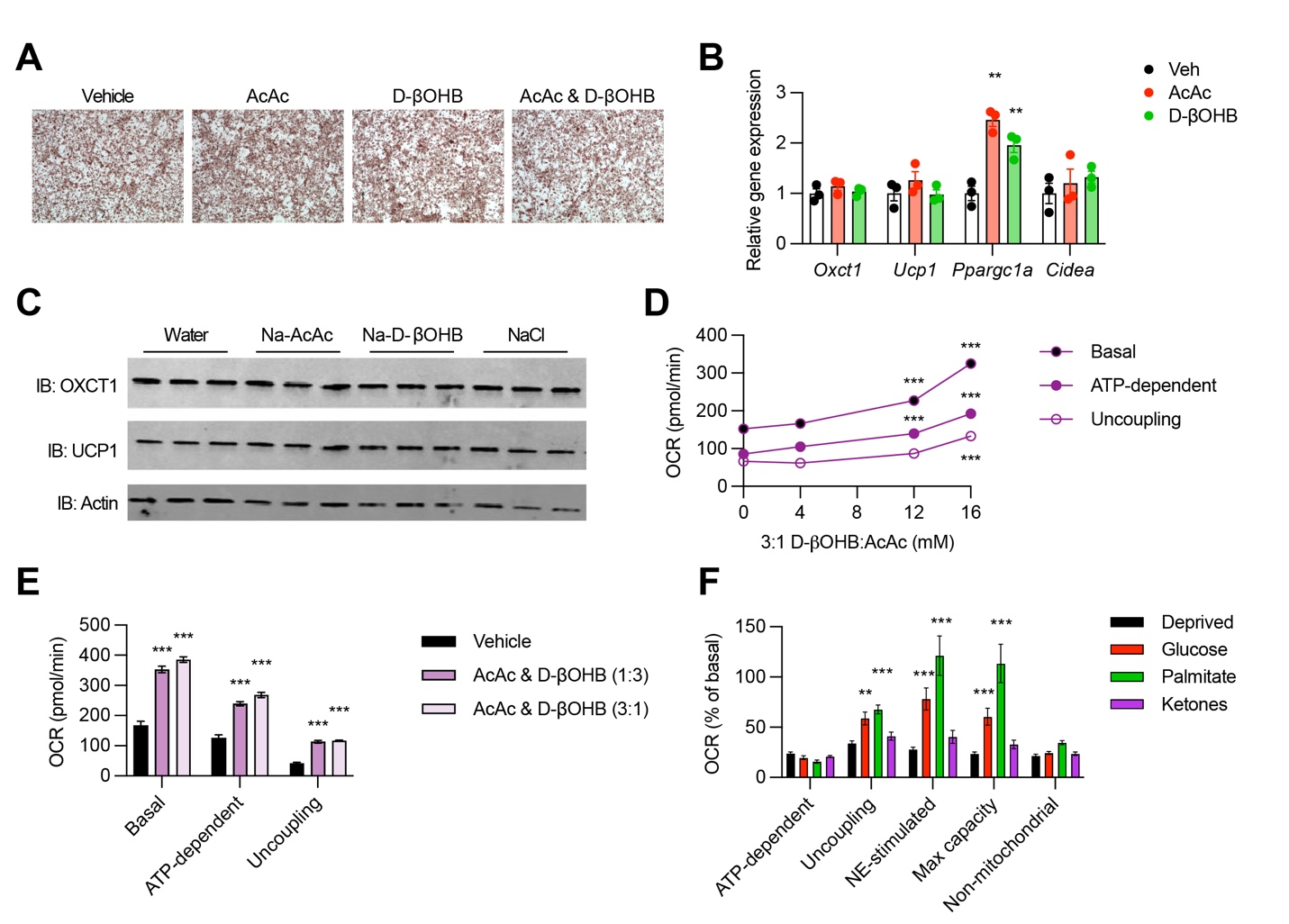

**Figure S3. Related to Figure 3. Ketone bodies promote oxygen consumption in brown adipocytes.**

(**A**) SVF cells from BAT were induced for adipogenesis with individual or mixed ketone bodies supplemented in the medium. Oil red O staining was performed.

(**B, C**) Primary brown adipocytes treated with AcAc or D-βOHB for 24h were subjected for RT-qPCR (B) and Western blotting (C).

(**D**) Basal, ATP-linked, and uncoupling respiration of primary brown adipocytes treated with varying concentrations of ketone bodies, analyzed by Seahorse respirometry.

(**E**) Seahorse analysis of primary brown adipocytes treated with 4 mM ketone bodies with reversing ratios of AcAc and D-βOHB.

(**F**) Relative OCR of primary brown adipocytes pretreated with glucose-deprived medium or supplemented with glucose, mixed ketones, or palmitate (Fig. 3E).

Data are represented as mean ± SEM. * P < 0.05, ** P < 0.01, *** P < 0.001 by one-way ANOVA.

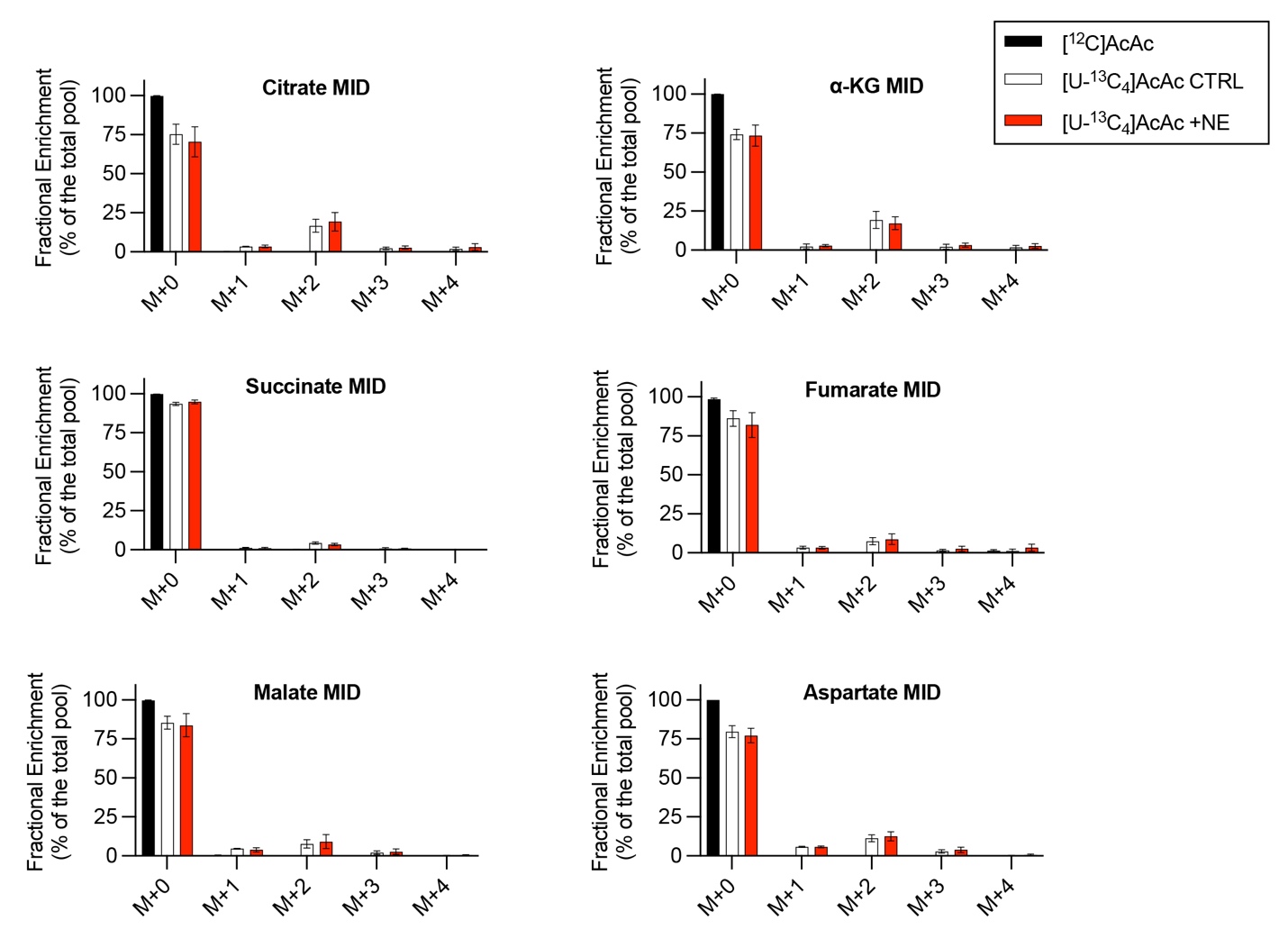

**Figure S4. Related to Figure 4. Enrichment of [U-^13^C_4_]AcAc to TCA intermediates.**

Mass isotopologue distribution (MID) of TCA cycle intermediates in primary brown adipocytes treated with 1 mM sodium [^12^C]AcAc, sodium [U-^13^C_4_]AcAc, or sodium [U-^13^C_4_]AcAc plus NE for 24 h.

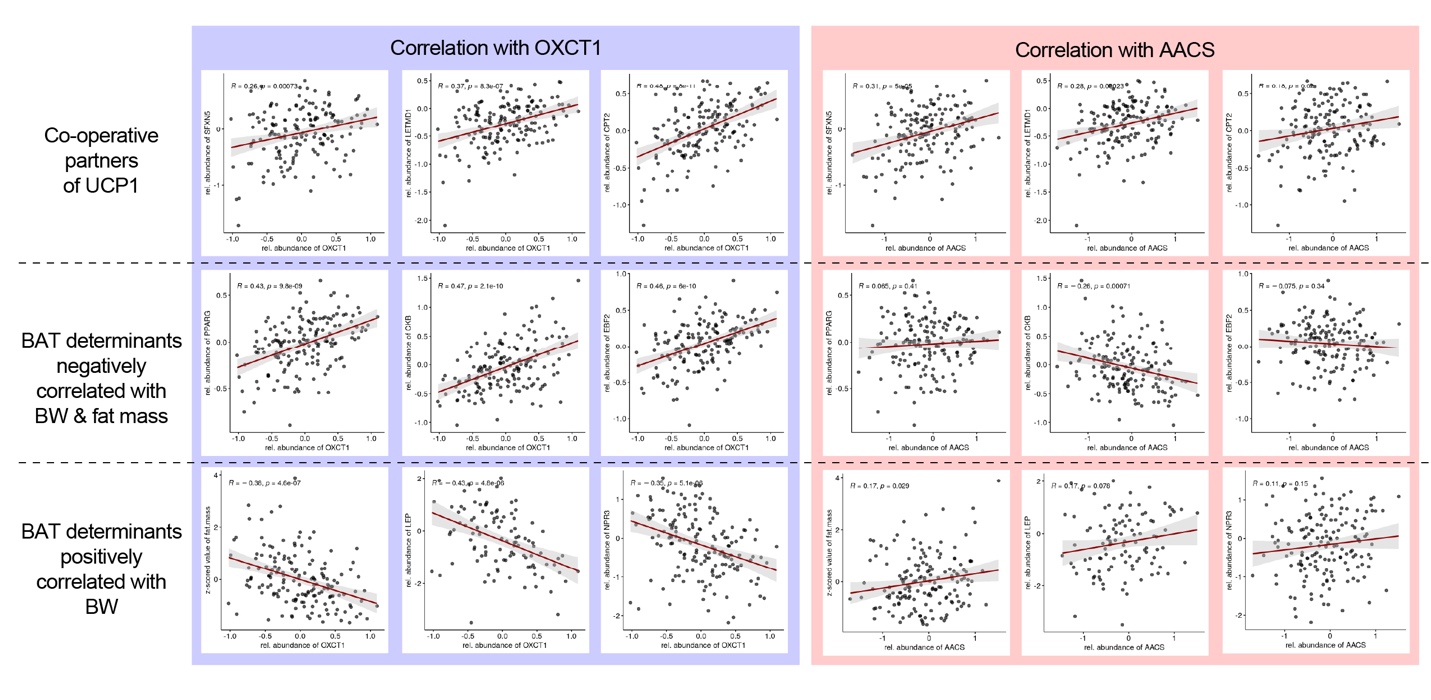

**Figure S5. Related to Figure 4 and 5.**

Regression analysis of OXCT1 (left) and AACS (right) with co-operative partners of UCP1 (SFXN5, LETMD1, and CPT2), BAT determinants negatively correlated with BW & fat mass (PPARG, CKB, and EBF2), and BAT determinants positively correlated with BW (fat mass, LEP, NPR3) in outbred mice.

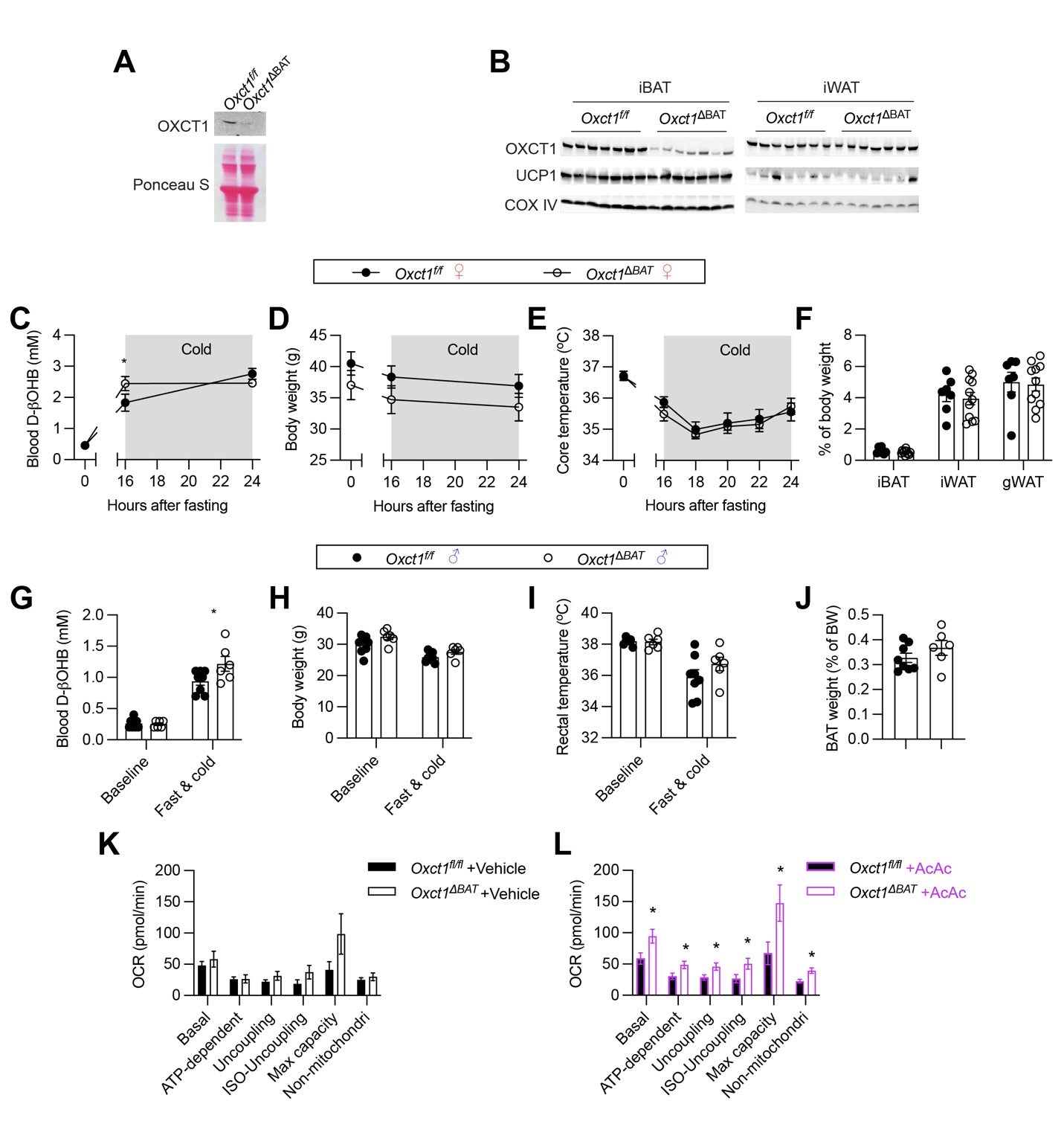

**Figure S6. Related to Figure 4. Phenotyping of *Oxct1^ΔBAT^* mice.**

(**A**) SVF and mature adipocytes were isolated from BAT of *Oxct1^f/f^* and *Oxct1^ΔBAT^* mice. OXCT1 expression was determined by Western Blotting.

(**B**) Western Blotting of OXCT1 and UCP1 in BAT and WAT from *Oxct1^f/f^* and *Oxct1^ΔBAT^* mice.

(**C-J**) Normal chow-fed female (C-F) and male (G-J) *Oxct1^f/f^* and *Oxct1^ΔBAT^* mice were challenged with fast and cold. Blood D-βOHB levels (C, G), body weight (D, H), rectal temperature (E, I), and tissue weight (F, J) were measured.

(**K, L**) Calculated OCR of primary brown adipocytes from *Oxct1^f/f^* and *Oxct1^ΔBAT^* mice in the absence (K) or presence (L) of AcAc. (Original Seahorse plot in Fig. 4H, I).

Data are represented as mean ± SEM. * P < 0.05 by two-way ANOVA (C, G) or two-tailed unpaired Student’s t-test (L).

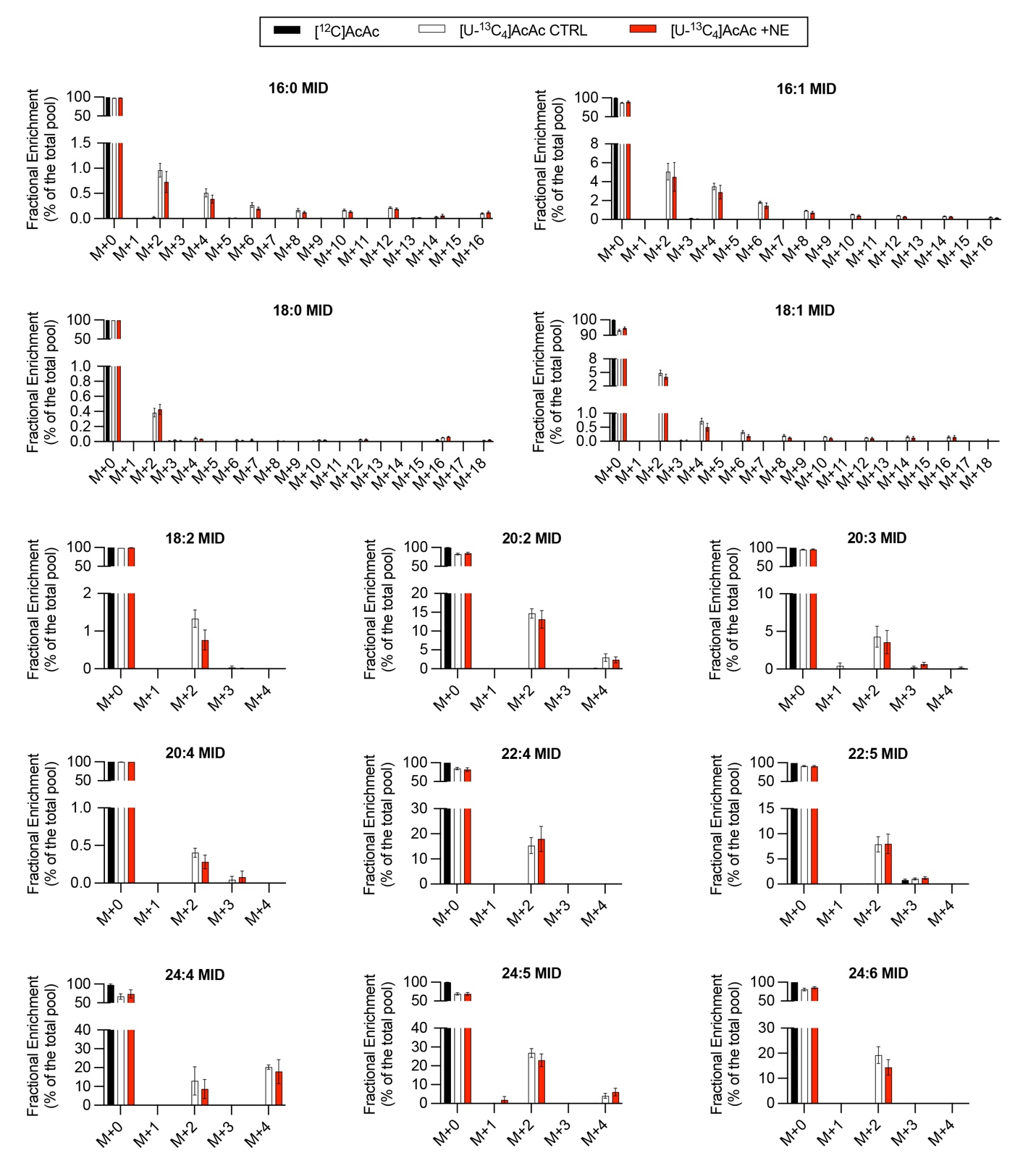

**Figure S7. Related to Figure 5.**

Enrichment of [U-^13^C_4_]AcAc to fatty acids in primary brown adipocytes treated with/without NE.

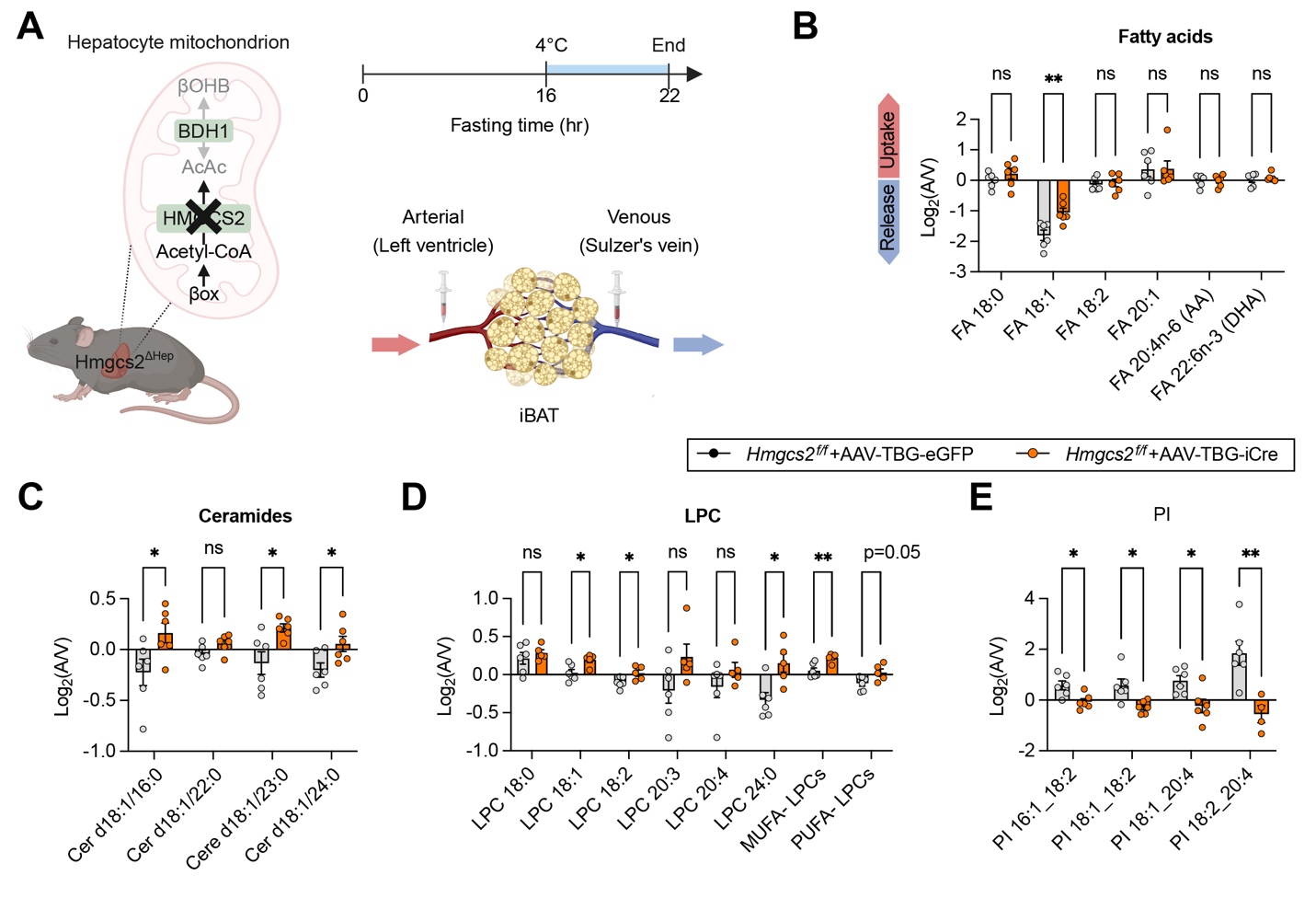

**Figure S8. Related to Figure 5. Metabolomics of AV samples from *Hmgcs2^ΔHep^* mice.**

(**A**) *Hmgcs2^ΔHep^* mice were subjected to 16h fasting followed with 8 h cold challenge. AV blood samples were collected for metabolics.

(**B**) Log2 values of A/V ratio showing net consumption (positive) and release (negative) of fatty acids (B), ceramides (C), LPC (D), and PI (I).

Data are represented as mean ± SEM. * P < 0.05, ** P < 0.01 by two-tailed unpaired Student’s t-test.

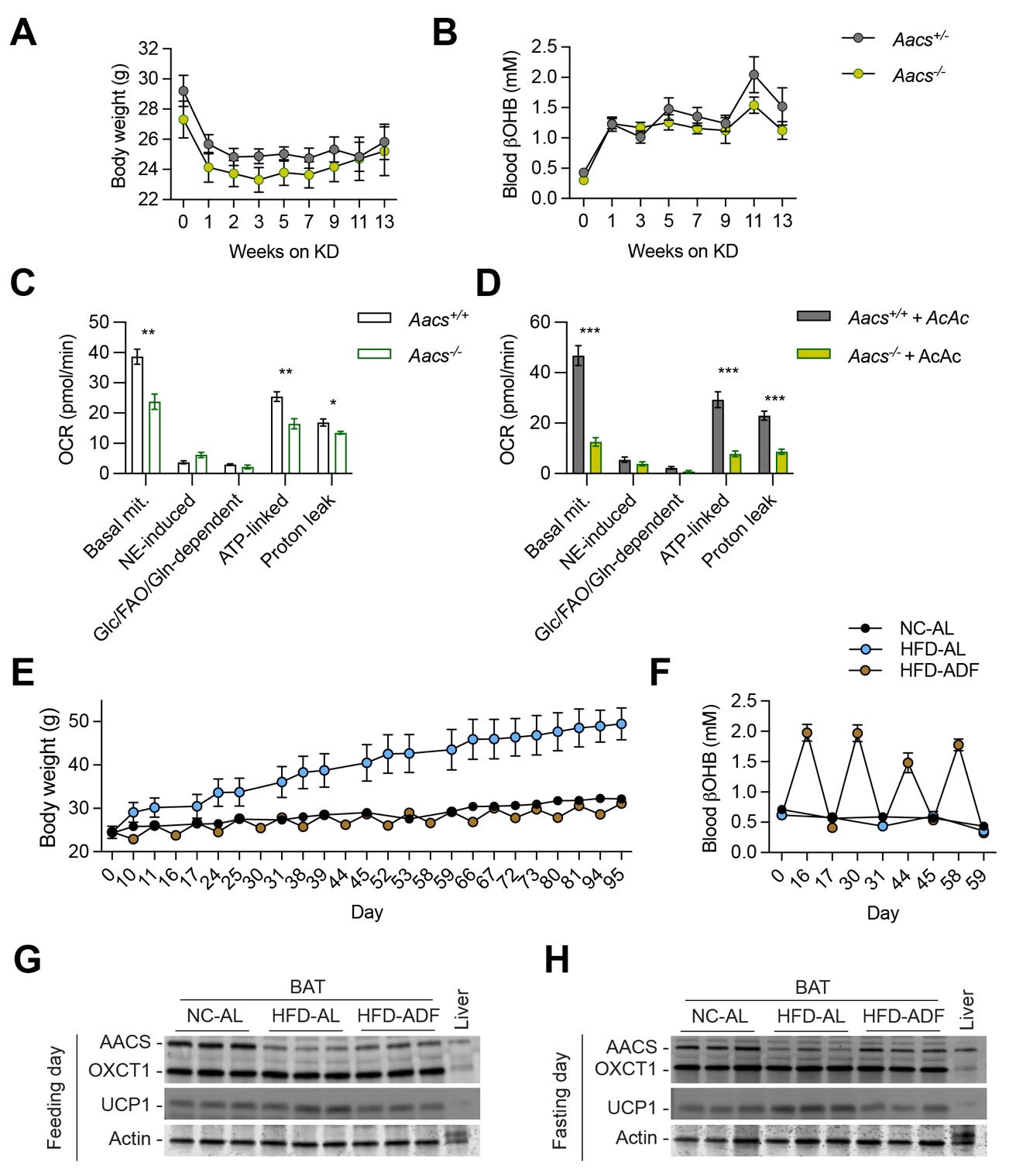

**Figure S9. Related to Figure 6. Phenotyping of *Aacs^-/-^* mice.**

(**A, B**) *Aacs*^+/+^ and *Aacs*^-/-^ mice were subjected to ketogenic diet feeding. Body weight (A) and blood D-βOHB levels (B) were shown.

(**C, D**) Calculated OCR values of primary brown adipocytes from *Aacs*^+/+^ and *Aacs*^-/-^ mice in assay medium containing 0 mM (C) or 3 mM (D) AcAc. Raw Seahorse plots in Figure 6C, D.

(**E-H**) C57BL/6J mice were fed with normal chow-ad libitum (NC-AL), HFD-AL, or HFD-alternate day fasting (ADF). Body weight (E) and blood D-βOHB levels (F) were determined on feeding and fasting days. Expression of AACS, OXCT1, and UCP1 were assessed by Western blotting in BAT tissues collected on feeding (G) or fasting (H) days.

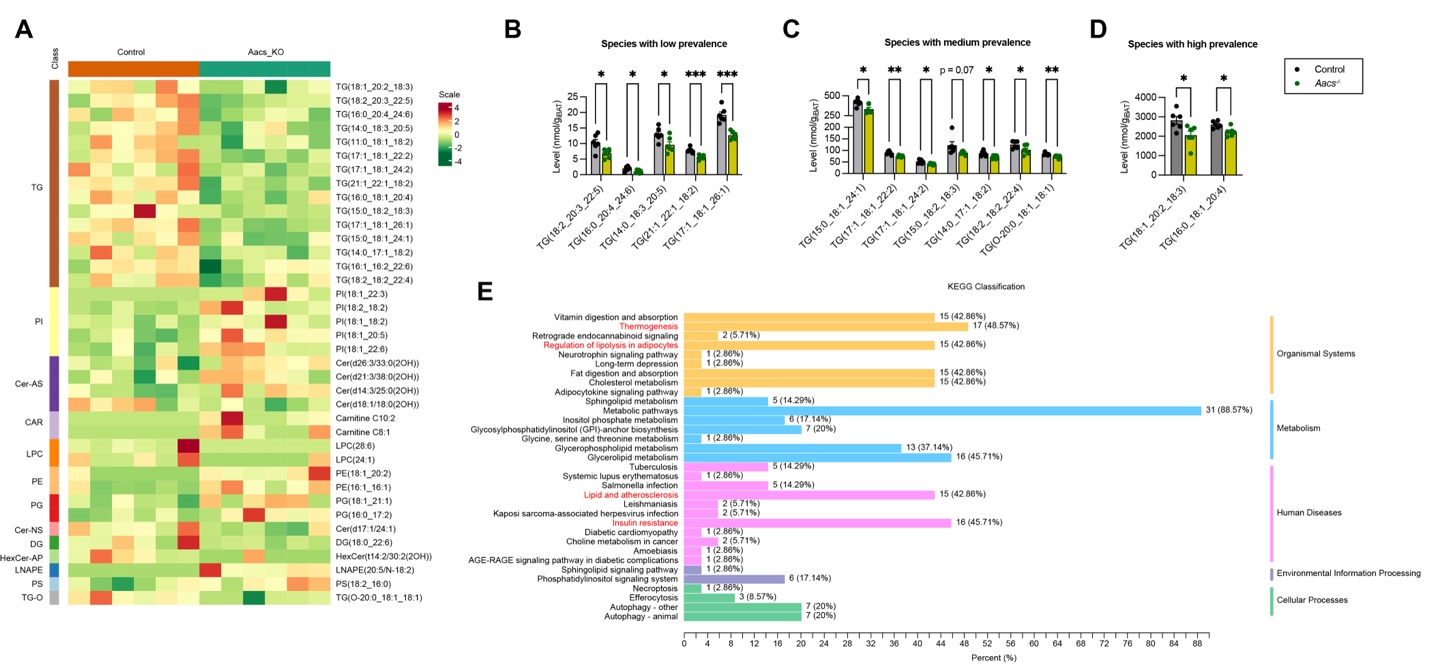

**Figure S10. Related to Figure 6. Lipidomics of BAT from *Aacs^-/-^* mice.**

(**A**) Heatmap of differential lipids in BAT between control and *Aacs*^-/-^ mice.

(**B-D**) Levels of TG species with low (B), medium (C), and high (D) prevalence in BAT of control and *Aacs*^-/-^ mice. Data are represented as mean ± SEM. * P < 0.05, ** P < 0.01, *** P < 0.001 by two-tailed unpaired Student’s t-test.

(**E**) KEGG pathway analysis of significantly differential lipids in BAT between control and *Aacs*^-/-^ mice.

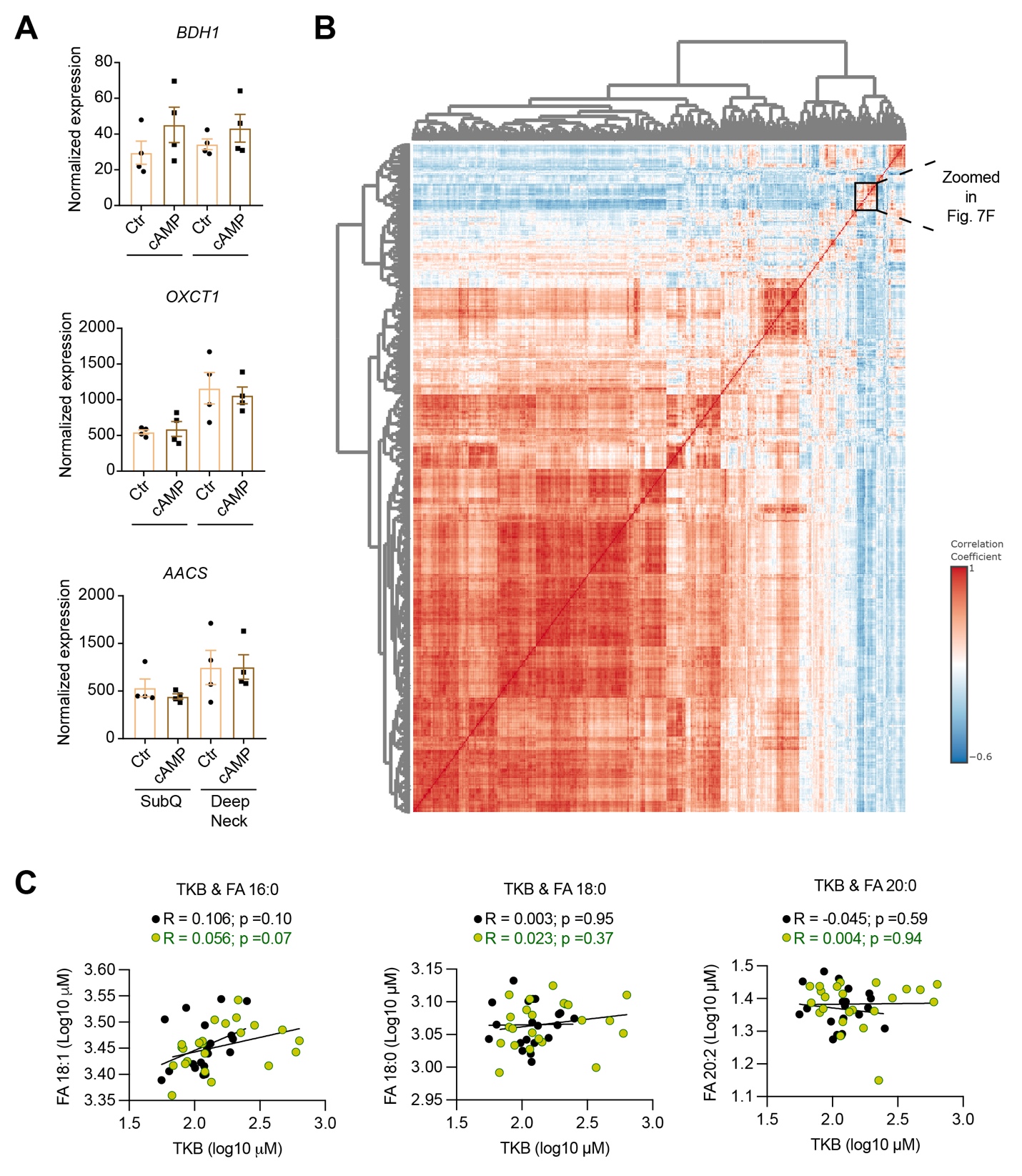

**Figure S11. Related to Figure 7. Regulation of BAT and lipids by ketone metabolism.**

(**A**) Preadipocytes from human subcutaneous (SubQ) WAT and deep neck BAT were differentiated into adipocytes and treated with or without cAMP. Expression of *BDH1*, *OXCT1*, and *AACS* genes were determined by RT-qPCR.

(**B**) Correlation of plasma lipid species from post-treatment UE and TRE patients.

(**C**) Regression analysis of post-treatment plasma AcAc level with saturated FAs in UE and TRE patients.
